# Molecular features underlying *Pseudomonas aeruginosa* persistence in human plasma

**DOI:** 10.1101/2021.12.21.473675

**Authors:** Manon Janet-Maitre, Stéphane Pont, Frerich Masson, Serena Sleiman, Julian Trouillon, Mylène Robert-Genthon, Benoît Gallet, Chantal Dumestre-Perard, Sylvie Elsen, Christine Moriscot, Bart Bardoel, Suzan Rooijakkers, François Cretin, Ina Attrée

**Author notes:** Laboratory of Pathogen-Host Interactions (LPHI), Université Montpellier, CNRS, UMR5235, 34095 Montpellier cedex 5, France. Institute of Molecular Systems Biology, ETH Zurich, Zurich, Switzerland.

## Abstract

*Pseudomonas aeruginosa*, an opportunistic Gram-negative pathogen, is a leading cause of bacteremia with a high mortality rate. We recently reported that *P. aeruginosa* forms a persister-like sub-population of evaders in human plasma and blood. However, the molecular mechanisms underlying the formation of evaders remained unknown. Here, using a gain-of-function genetic screen, we examined the molecular determinants of *P. aeruginosa* persistence in plasma. We found that, among other factors, ATP and biotin availability greatly influence bacterial survival in plasma; mutants in *pur* and *bio* genes display higher tolerance and persistence, respectively. Electron microscopy combined with energy-dispersive X-ray spectroscopy (EDX) revealed the formation of polyphosphate granules upon incubation in plasma in several clinical strains, implying the bacterial response to a low-energy stress signal. Indeed, mutants with transposon insertions in *ppk* genes were eliminated in the plasma. Analysis of several steps of the complement cascade and exposure to an outer-membrane-impermeable drug, nisin, suggested that the mutants impede membrane attack complex (MAC) activity *per se*. Through this study, we shed light on *P. aeruginosa* response to the plasma conditions and discovered the multifactorial origin of bacterial resilience to MAC that provides a comprehensive picture of the complex interplay between *P. aeruginosa* and the human complement system.

**Author summary:** Persistence of bacterial pathogens is a main cause of treatment failure and establishment of chronic bacterial infection. Despite innate immune responses, some bacteria may persist in human blood and plasma. Here we used a genome-wide screen to investigate the molecular determinants influencing *Pseudomonas aeruginosa* persistence in human plasma facing the complement system. Alongside a multifactorial strategy, we found intracellular levels of ATP and biotin to significantly influence bacterial capacity to deal with membrane attack complex (MAC)-dependent killing. These results underline the need to understand the complex interplay between bacterial pathogens and the human immune system when seeking to develop efficient antibacterial strategies.

## Introduction

Human bacterial pathogens employ sophisticated strategies to escape control by the immune system, and to resist or tolerate antibiotic treatments. Bacterial persistence toward antibiotics is a major obstacle in the treatment of life-threatening infections and is frequently associated with the establishment of chronic infections, as it allows the survival of a subpopulation of bacteria that are tolerant to what should be lethal antibiotic stress (Bartell et al., 2020; Fauvart et al., 2011). Unlike antibiotic resistance, persistence is a non-heritable, fully reversible trait characterized by a biphasic killing curve – with rapid killing of the bulk population and survival of a subpopulation over a long period of time. Among the factors hypothesized to lead to antibiotic persistence (Balaban et al., 2019), the most cited are low intracellular ATP levels (Manuse et al., 2021; Shan et al., 2017) and production of the “alarmone” signaling molecule guanosine (penta) tetraphosphate, (p)ppGpp (Korch et al., 2003; Pacios et al., 2020). Following external antibiotic stress, stochastic or elicited activation of the stringent response leads to accumulation of (p)ppGpp. This molecule in turn activates toxin-antitoxin modules which trigger a regulatory cascade leading to degradation of the antitoxins, ultimately resulting in inhibition of cell growth (Gaca et al., 2015; Harms et al., 2016). The host immune system also elicits persistence in a number of bacterial species. For example, acidification of macrophage vacuoles after *Salmonella* internalization induces persister cell formation and could contribute to establishing a bacterial reservoir for infection relapse (Helaine et al., 2014). In a similar manner, exposure to human serum triggers the formation of antibiotic persisters as well as so-called “viable but non-culturable” forms of *Vibrio vulnificus* (Ayrapetyan et al., 2015). Putrinš *et al.* (Putrinš et al., 2015) reported that phenotypic heterogeneity in *Escherichia coli* could serve as a means to evade serum-mediated and antibiotic-induced killing. In a previous study (Pont et al., 2020), we demonstrated that a persister-like sub-population of *Pseudomonas aeruginosa* evaders forms when generally sensitive bacteria are incubated in human blood or human plasma. In this article, we refer to resistance, tolerance, and persistence as defined in a context of exposure to antibiotics (Balaban et al., 2019). Briefly, resistance is due to a heritable resistance factor allowing survival in the face of higher stress conditions. In contrast to resistance, tolerance is a phenomenon that allows survival of the population despite an otherwise lethal stress for a longer period of time, but without deploying any resistance mechanism. The third term, persistence, is the capacity of a tolerant sub-population to survive whereas the rest of the population is eliminated. Finally, we used the term resilience to describe several of these phenomena.

*P. aeruginosa* is a leading nosocomial pathogen that is extremely difficult to eradicate due to its high adaptability to a range of environments and its intrinsic and acquired antibiotic resistance (Moradali et al., 2017). Bloodstream infections (BSI) due to *P. aeruginosa* strains hold the record for highest mortality rate, at about 40% (Kang et al., 2003; Vitkauskienė et al., 2010). The complement system (CS) is the major immune component in human blood responsible for *P. aeruginosa* killing, and it is generally efficient against sensitive strains. However, a tiny fraction of the initial bacterial population can persist despite prolonged incubation in blood or plasma (Pont et al., 2020). The kinetics of *P. aeruginosa* killing by human plasma displays a classical persister-type biphasic profile (Balaban et al., 2019), with rapid elimination of more than 99.9% of the population, reaching a plateau at an average of 0.1% survival after 2 h. In our previous study, we found that, out of 11 BSI isolates tested, 3 were plasma-resistant, 3 displayed a tolerant phenotype, and 5 were able to form evaders. These proportions suggest that evasion may represent a common strategy used by *P. aeruginosa* to escape membrane attack complex (MAC)-mediated bactericidal activity in plasma. The plasma evaders were distinct from antibiotic persisters since a stationary phase population did not give rise to a higher population of evaders, as reported for antibiotic persisters (Keren et al., 2004).

For this study, we employed a genome-wide approach to obtain first insights into the molecular mechanism(s) controlling *P. aeruginosa* persistence in human plasma. A screen based on transposon-insertion sequencing (Tn-seq) was applied to a recent clinical isolate that was sensitive to plasma, but produced between 0.01 and 0.1% of evaders after 6 h in plasma. We hypothesized that transposon mutants for which an increased proportion of evaders was detected or mutants with increased overall plasma tolerance/resistance could reveal novel determinants of bacterial resilience to plasma and pathogen immune escape. In addition to *P. aeruginosa* factors known to be involved in plasma resistance, such as O-specific antigen (OSA) length and exopolysaccharides, we identified an uncharacterized small periplasmic protein that increases complement tolerance up to 100-fold. Our screen and the associated phenotypic characterization of mutants also revealed that biotin and ATP levels modulate pathogen sensitivity to MAC-induced lysis.

## Results

### Gain-of-function screen in plasma

In our previous study, we found that the *Pseudomonas aeruginosa* isolate IHMA879472/AZPAE15042 (IHMA87) persisted in human blood and plasma. Although the overall population was sensitive to complement killing, with 99.9% of bacteria eliminated within 2 hours, approximately 0.1% of evaders survived prolonged incubation in plasma (Pont et al., 2020).

To search for mutants generating higher number of evaders using a genome-wide approach, we constructed a transposon library of about 300,000 mutants in IHMA87 using the Himar-1 transposon. The transposon library was challenged in native human plasma (output) or heat inactivated plasma (HIP, input) for three hours before harvesting. Bacterial survival was expressed as colony forming unit (CFU) counts for the whole library before and after the challenge. This analysis revealed that overall survival was increased by about 2-log in the mutant library compared to the parental strain. Thus, the screen effectively selects for this type of mutants (Fig. 1A). Resistant, hyper-evader, and tolerant phenotypes can then be differentiated based on killing kinetics during long incubation in plasma.

**Fig 1.**
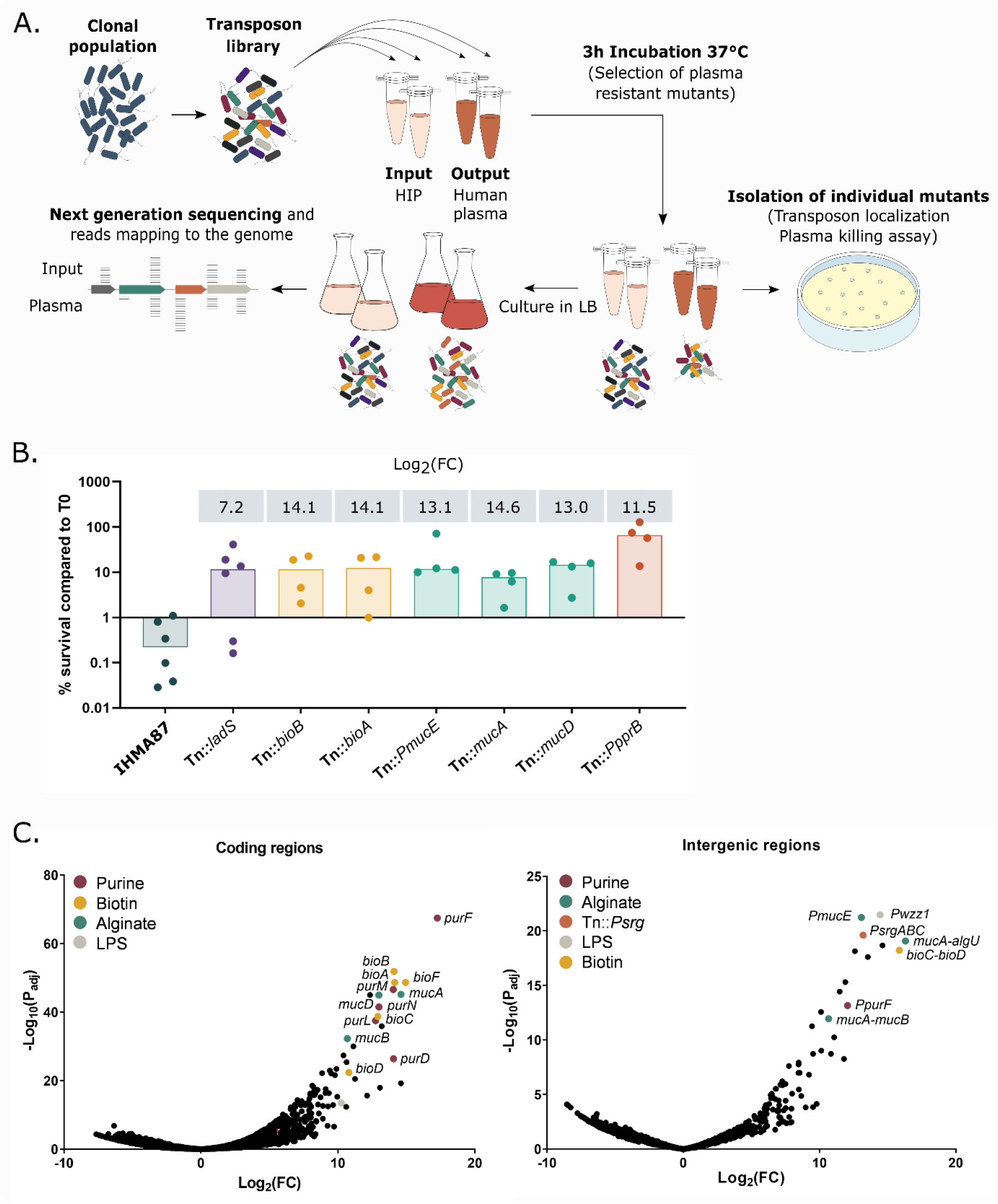
Gain-of-function Tn-seq reveals common and novel pathways contributing to *P. aeruginosa* survival in plasma. **A.** Schematic representation of the screening method used for this study. A transposon-insertion mutant library was generated in the plasma-sensitive clinical isolate *P. aeruginosa* IHMA87 and exposed either to human plasma or to HIP (input) for 3 h. Aliquots were plated on LB and individual transposon mutants were stored for further analysis. **B.** Survival of isolated transposon mutants following incubation for 3 h in plasma. Survival rates were calculated based on CFU measurements. The dataset was log-transformed, and rates for all mutants were significantly different from rates for the wild-type strain (*p*-value < 0.005). Log_2_(Fold-Change(FC)) obtained from the Tn-seq analysis for each gene or intergenic region is indicated above the histogram. **C.** Bioinformatics analyses presented as Volcano plots. Insertions in coding regions (left) and intergenic regions (right) were analyzed separately. Significant hits in genes involved in the same pathway are shown in the same color.

Seven mutants of interest were isolated during the screen, and their survival was assayed in plasma killing assays. Survival rates for these mutants were between 10- and 200-fold higher than those of the parental strain (Fig. 1B), thus validating the screen. Random colony-picking failed to identify any LPS-related mutants. However, the screen did confirm the importance of LPS and more specifically OSA biosynthesis in the bacteria’s interaction with plasma (Goldberg and Pler, 1996; Kintz et al., 2008; Osawa et al., 2013; Priebe et al., 2004).

Sequencing data was analyzed both for insertions in coding regions and in intergenic regions, as the transposon included an outward *tac* promoter, which could modulate the expression of neighboring genes. In both analyses, transposon insertions strongly increasing bacterial survival were associated with three main pathways: biosynthesis of purine and biotin; and biosynthesis, production, and regulation of the exopolysaccharide alginate (Fig. 1C and Table 1, full results in Table 1S).

**Table 1.** Mutants enriched in plasma compared to HIP carry insertions in three main pathways. * IHMA87 gene identity (ID) (NCBI accession numbers CP041354 and CP04135), for the chromosome and plasmid sequences respectively (Trouillon et al., 2020) are indicated with PAO1 equivalents ** where applicable. IG is added when insertions were detected in the <u>i</u>nter<u>g</u>enic region between two genes and the intergenic region is attributed to the upstream gene, independently on the genes orientation. ^#^ For each gene, the Log_2_(FC) in plasma compared to HIP (input) condition is indicated together with the corresponding adjusted *p*-value (P_adj_ ^##^), obtained by the DeSeq2 program. ^$^ The genes are sorted according to their predicted pathway taken from the KEGG database and the *Pseudomonas* genome database (Kanehisa et al., 2008; Winsor et al., 2011).

| IHMA87 ID * | PAO1 ID ** | gene/prediction | Log<sub>2</sub>(FC) # | *P*<sub>adj</sub><sup>##</sup> |
| --- | --- | --- | --- | --- |
| **Purine biosynthetic pathway \$** | | | | |
| *IHMA87_01837* | *PA3108* | *purF* | 17.26 | 3.64E-68 |
| *IHMA87_05343* | *PA4855* | *purD* | 14.06 | 3.85E-27 |
| *IHMA87_04292* | *PA0944* | *purN* | 12.99 | 3.14E-42 |
| *IHMA87_04291* | *PA0945* | *purM* | 14.04 | 2.51E-47 |
| *IHMA87_01191* | *PA3763* | *purL* | 12.75 | 3.02E-38 |
| *IHMA87_04214* | *PA1013* | *purC* | 7.67 | 1.23E-5 |
| *IHMA87_02487* | *PA2629* | *purB* | 5.35 | 7.23E-4 |
| **Biotin biosynthetic pathway** |  |  |  |  |
| *IHMA87_00400* | *PA0420* | *bioA* | 14.15 | 2.29E-49 |
| *IHMA87_00477* | *PA0501* | *bioF* | 14.94 | 2.29E-49 |
| *IHMA87_00476* | *PA0500* | *bioB* | 14.10 | 1.39E-52 |
| *IHMA87_00479* | *PA0503* | *bioC* | 12.92 | 1.90E-39 |
| *IHMA87_00480* | *PA0504* | *bioD* | 10.80 | 4.23E-23 |
| **Alginate biosynthetic/regulation genes** |  |  |  |  |
| *IHMA87_04494* | *PA0762* | *algU* | 11.25 | 3.01E-21 |
| *IHMA87_04493* | *PA0763* | *mucA* | 14.60 | 6.76E-46 |
| *IHMA87_04492* | *PA0764* | *mucB* | 10.69 | 5.56E-33 |
| *IHMA87_04490* | *PA0766* | *mucD* | 13.00 | 1.10E-45 |
| *IHMA87_00895* | *PA4033* | *P<sub>mucE</sub>* | 13.07 | 5.88 E-22 |
| *IHMA87_05847* | *PA5322* | *algC* | 2.66 | 4.76 E-2 |
| **Other top hits** |  |  |  |  |
| *IHMA87_02894* | NA | -/TraX family protein | 14.60 | 6.18E-20 |
| *IHMA87_02666* | *PA2484* | -/transcriptional regulator | 13.20 | 1.33E-36 |
| *IHMA87_01574_IG* | *PA3368_IG* | *P<sub>srg</sub>* | 13.19 | 2.50E-20 |
| *IHMA87_01786_IG* | *PA3161_IG* | *P<sub>wzz1</sub>* | 12.59 | 7.33E-19 |
| ***IHMA87_06317*** | NA | - | 12.34 | 1.02E-45 |
| ***IHMA87_02554*** | *PA2573* | -/chemotaxis<br>transducer | 11.13 | 1.02E-30 |
| ***IHMA87_06033*** | *PA5490* | *cc4* | 10.63 | 4.10E-26 |
| ***IHMA87_04274*** |  | *slyX* | 10.60 | 4.67E-13 |
| ***IHMA87_04133*** | *PA1078* | *flgC* | 10.40 | 4.59E-28 |
| ***IHMA87_04779*** | *PA4457* | *kdsD*/arabinose-5-phosphate<br>isomerase | 10.24 | 3.15E-14 |
| ***IHMA87_00319*** | *PA0341* | *lgt* | 9.95 | 5.21E-16 |
| ***IHMA87_00896*** | *PA4032* | -/probable<br>response regulator | 9.89 | 3.67E-24 |
| ***IHMA87_03447*** | *PA1758* | *pabB* | 9.79 | 2.55E-22 |
| ***IHMA87_05840*** | *PA5315* | *rpmG* | 9.71 | 2.60E-07 |
| ***IHMA87_03022*** | *PA2122* | -/lipid biosynthetic<br>process | 9.65 | 1.27E-13 |
| ***IHMA87_01755*** | NA | - | 9.51 | 7.37E-23 |
| ***IHMA87_04617*** | *PA4315* | *mvaT* | 9.41 | 1.21E-23 |
| ***IHMA87_02603*** | NA | - | 9.34 | 4.46E-17 |
| ***IHMA87_02570*** | NA | - | 9.28 | 3.29E-13 |
| ***IHMA87_05344*** | *PA4856* | *retS* | 9.11 | 2.94E-18 |

### Analysis of isolated mutants

Two isolated displayed hyper-mucoid phenotypes – reflecting alginate overproduction – associated with insertions within *mucD* and in the intergenic region upstream of *mucA*. These two genes are negative regulators of alginate synthesis (reviewed in Damron and Goldberg, 2012). We also isolated a mutant in which the transposon was inserted into the intergenic region upstream of *mucE*, encoding a positive regulator of alginate production (Qiu et al., 2007; Yin et al., 2013). The orientation of the transposon suggested that this mutant could overexpress *mucE*, also resulting in a mucoid phenotype. Alginate overproduction may interfere with the CS by decreasing C3b-dependent opsonization and thus preventing CS activation upon acetylation (Pedersen et al., 1990; Pier et al., 2001).

An alternative mechanism was revealed by a transposon insertion in *ladS*, encoding a histidine kinase that positively regulates the GacS/GacA-RsmA pathway (Broder et al., 2017; Ventre et al., 2006), which was also associated with increased mutant survival. The downstream effector of this pathway is as yet unknown. Interestingly, bioinformatics analysis of the sequencing data revealed significant enrichment of transposon insertions in *retS*, which codes for a LadS antagonist (Table 1). This effect is of interest as Psl improved bacterial survival of a mucoid strain in serum (Jones and Wozniak, 2017). The role of Psl in complement resistance might be strain-dependent, as assays using the reference strain PAO1 initially showed that Psl reduces bacterial opsonization without affecting C9 insertion or bacterial survival in serum (Mishra et al., 2012).

Survival in plasma was also increased by insertion of the transposon in the promoter of *pprB* encoding the response regulator acting on type IVb pili/Tad, BapA adhesin and CupE fimbriae expression (de Bentzmann et al., 2012; Giraud et al., 2011). As shown in *Vibrio cholerae* and *Neisseria gonorrhoeae*, type IV pili contribute to serum resistance by recruitment of the host negative complement regulator C4BP (Blom et al., 2001; Chiang et al., 1995). Our data thus suggest that either type IVb pili or CupE fimbriae could be also involved in *P. aeruginosa* plasma resilience, but the mechanisms involved need to be explored further.

Finally, in two isolated mutants showing increased survival, the transposon was inserted into *bioA* and *bioB*, encoding enzymes involved in biotin biosynthesis. To our knowledge, these genes have not previously been linked to resilience to plasma or complement.

Together, those results confirm the role of LPS, alginates, and Psl in resistance to the CS, validating the Tn-seq approach. In addition, several novel determinants worth exploring were identified based on mutants showing enhanced survival (>9 Log_2_(FC)) following transposon insertion in uncharacterized genes (Table 1).

### Expression of the three-gene operon *srgABC* improves survival in plasma

Sanger sequencing of one mutant isolated during the screen that was significantly enriched in plasma (Log_2_(FC) of 13.2; Fig. 2A) revealed transposon insertion within the intergenic region between an uncharacterized predicted three-gene operon *IHMA87_01573-IHMA87_01571 (PA3369-PA3371* in PAO1) and *panM* (*IHMA87_01574/PA3368*), a probable acetyltransferase (Fig. 2B). The mutant isolated displayed a tolerant phenotype over a 6-h period (Fig. S1), with survival increased 100-fold compared to the parental strain after 3 h (Fig. 2C).

**Fig 2.**
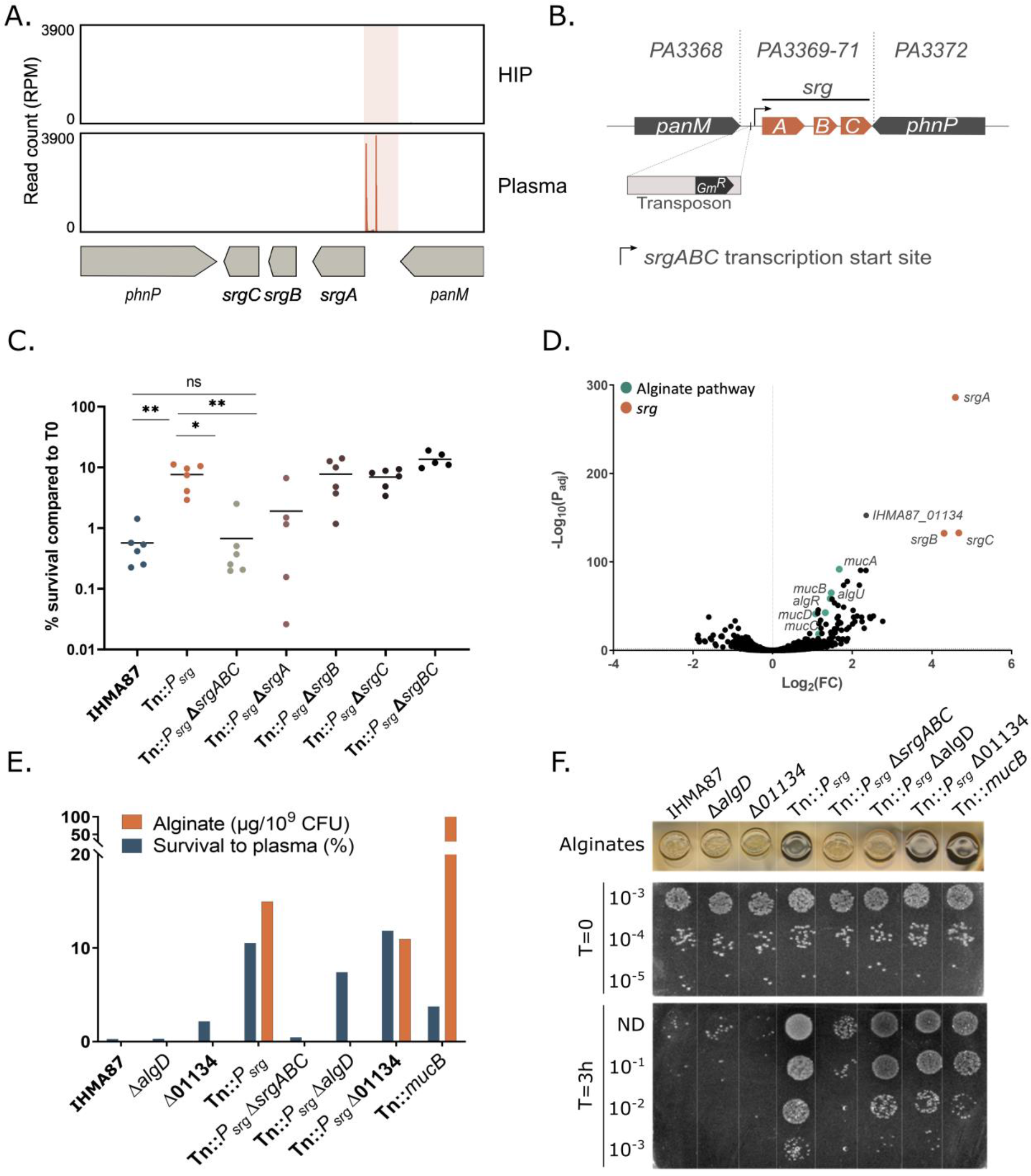
Overexpression of *srgA* increases survival 100-fold. **A.** Zoom in on the Tn-seq profiles in the region surrounding the *srg* operon, showing the number of normalized reads in input (HIP) and output (plasma). **B.** Schematic representation of the predicted *srgABC* operon and position of the transposon in the isolated Tn∷*P_srg_* mutant, as revealed by Sanger sequencing. Corresponding genes from PAO1 are indicated. Note that the transposon is inserted 13 bp upstream of the predicted transcriptional start site. **C.** Role of individual *srg* genes in the plasma resilience phenotype of the Tn∷*P_srg_* strain. Deletion of *srgA* restores sensitivity to plasma. **D.** Differential gene expression between Tn∷*P_srg_* and the parental IHMA87 strain represented in a volcano plot showing overexpression of *srgABC* and alginate-related genes. RNA was extracted from cultures grown on LB, and whole transcriptomes were determined by an RNA-seq pipeline **E-F.** Survival in plasma and alginate production for *P. aeruginosa* strains. Alginate synthesis was visualized based on colony morphology and quantified by carbazole assay, as described in Materials and Methods. Alginate overproduction appears as a darker bacterial spot (**F**). The data shown are from one representative experiment, performed in biological triplicates.

The effect of the insertion was impossible to assess based on the position of the transposon and the RT-qPCR experiments. In addition, upon exposure to plasma, survival of strains bearing deletions in *panM*, or in *IHMA87_01573-IHMA87_01571* in the IHMA87 background was identical to survival of the wild-type strain (data not shown). Therefore, to get an idea of the overall transposon-induced changes in the mutant, we determined its transcriptomic profile and compared it to the parental strain in LB. RNA-seq data (Table 2S) revealed overexpression (about 20-fold compared to the parental strain) of *IHMA87_01573-IHMA87_01571*, named *srgABC* for serum resistance genes. Based on this result, the transposon-insertion mutant will hereafter be referred to as Tn∷*P_srg_*. In total, many genes were differentially expressed between Tn∷*P_srg_* and the wild-type strain (Table 2S), including *algU* – encoding the alternative sigma factor AlgU – and five other alginate genes (*algR* and *mucABCD* operon, Fig. 2D). In line with the observed upregulation of *algU*, Tn∷*P_srg_* colonies displayed a characteristic mucoid phenotype on solid medium.

The *srg* operon encodes three small putative proteins: SrgA (9.9 kDa), SrgB (5.3 kDa) and SrgC (6.5 kDa). Structural predictions indicate SrgA to be a periplasmic or secreted protein with a signal peptide cleavage site at position Gly27. Both SrgB and SrgC are predicted to be membrane proteins with one and two transmembrane alpha-helices, respectively. All three *srg* gene products are highly conserved (98-100% amino acid sequence identity) over the 232 complete *P. aeruginosa* genomes available in the *Pseudomonas* genome database (Winsor et al., 2016).

The individual contributions of the *srg* genes to *P. aeruginosa* tolerance to plasma were evaluated by testing single, double and triple deletion mutants directly in the Tn∷*P_srg_* background strain. Plasma killing assays performed with these deletion mutants showed that SrgA was required and sufficient for increased survival in plasma (Fig. 2C).

### Purine and biotin pathway deficiencies lead to increased persistence

We then focused on mutants where the transposon was present in genes and intergenic regions linked to biotin and purine biosynthesis. Mutants with insertions in the *bioBFHCD* operon and in the *bioA* gene – covering all steps of biotin biosynthesis – were significantly overrepresented following plasma challenge (Table 1, Fig. 3A top). This result suggests that lack of biotin promotes survival in plasma. This finding was unexpected as the biotin pathway has been shown to be essential for survival in human serum for several pathogens, including *Klebsiella pneumoniae* and *Mycobacterium tuberculosis* (Carfrae et al., 2020; Weber et al., 2020; Woong Park et al., 2011).

**Fig 3.**
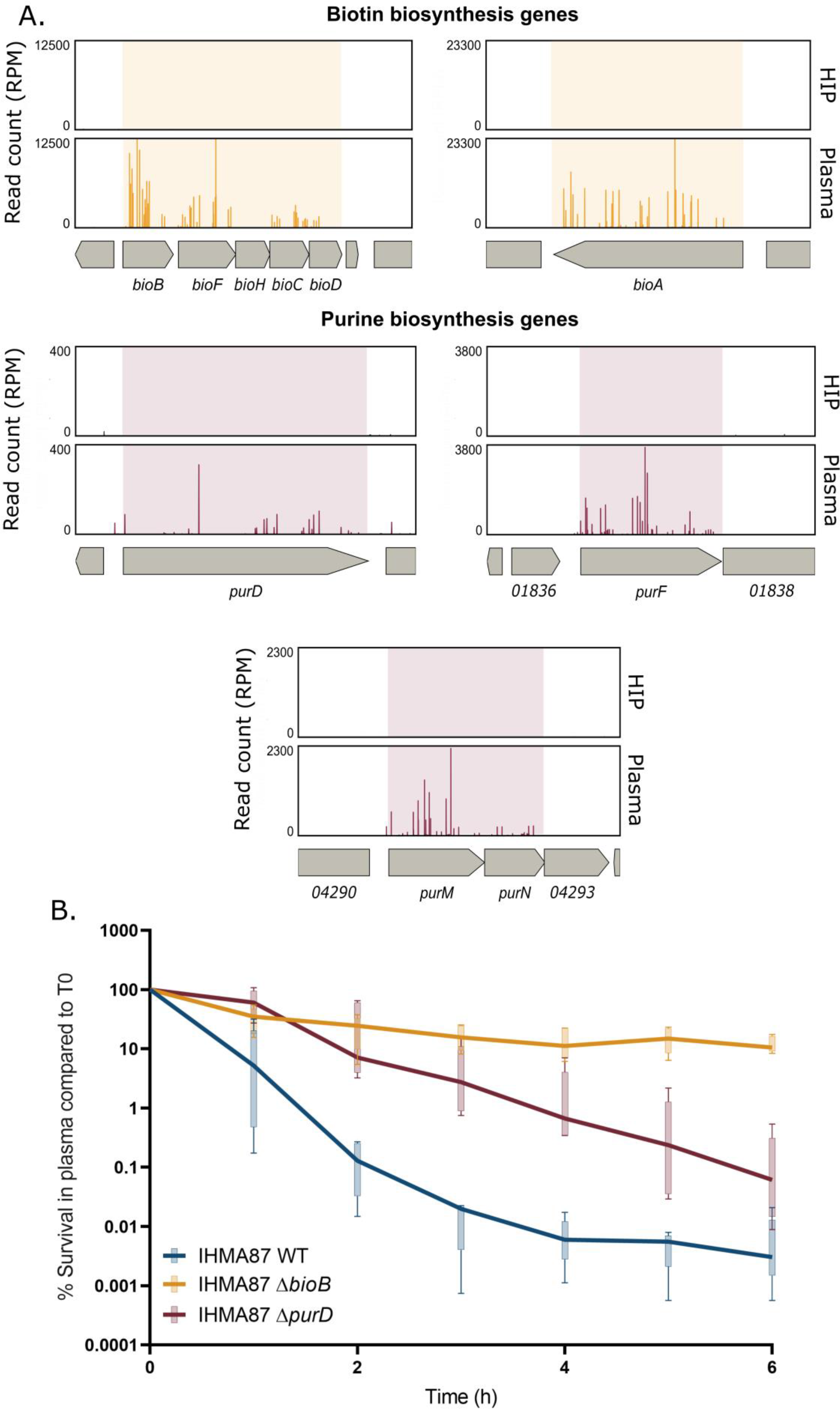
Inactivation of biotin and purine biosynthetic pathways increases survival rates of IHMA87 in plasma. **A.** Zoom in on Tn-seq profiles of *bio* and *pur* genes and operons, showing normalized numbers of reads in input (HIP) and output (plasma) samples. **B.** IHMA87 wild-type strain, Δ*bioB*, and Δ*purD* mutants’ survival kinetics in plasma over 6 h incubation, as measured by CFU counting (n=5). Note the biphasic killing curves for the parental strain and Δ*bioB* mutant, indicating increased persistence of Δ*bioB.* The Δ*purD* mutant displayed increased tolerance compared to the parental strain.

To confirm this result, in addition to the mutants isolated from the screen (Tn∷*bioA* and Tn∷*bioB*), we engineered a *bioB* deletion mutant (Δ*bioB*) and examined its behavior in plasma over a 6-h challenge (Fig. 3B). The Δ*bioB* mutant survived significantly better than the wild-type strain in plasma (over 10% vs. about 0.01% survival, respectively). Moreover, in plasma, IHMA87Δ*bioB* displayed a biphasic killing curve, implying that the levels of biotin determine the persistence levels.

*De novo* purine biosynthesis, together with pyrimidine production, are essential for bacterial nucleotide metabolism and important for bacterial growth and survival in plasma and serum (Andersen-Civil et al., 2018; Poulsen et al., 2019; Samant et al., 2008; Zhu et al., 2021). Like biotin synthesis, in our model, survival in plasma was clearly enhanced for bacteria bearing mutations in the purine biosynthetic pathway (Fig. 3A bottom). Transposon-insertion mutants in three main operons encoding enzymes involved in purine biogenesis were overrepresented in the screen. In particular, mutants in *purF*, encoding an amido-phospho-ribosyltransferase, were significantly enriched with a Log_2_(FC) of 17. To further study the link between purine biosynthesis and increased plasma resistance, we designed a *purD* deletion mutant (Δ*purD*) and studied its killing kinetics over 6 h in plasma. In accordance with Hill *et al*. in an antibiotic context (Hill et al., 2021), the Δ*purD* mutant displayed a tolerant phenotype in plasma (Fig. 3B). Interestingly, no similar level of gene enrichment was found for the pyrimidine pathway suggesting that the specific product of the purine pathway, rather than the nucleotide metabolism as a whole, contributed to increased survival in human plasma.

### ATP levels influence *P. aeruginosa* tolerance to plasma

Recent studies showed that low intracellular ATP concentration resulted in increased antibiotic persistence (Manuse et al., 2021; Shan et al., 2017). Along the same line, bacteria harvested at the stationary growth phase have lower levels of intracellular ATP and form a higher proportion of persisters when treated with antibiotics (Conlon et al., 2016). In our hands, the growth phase did not contribute significantly to the proportion of evaders, suggesting that the initial metabolic status of bacteria does not significantly contribute to persistence (Pont et al., 2020). However, because ATP is a final product of the purine pathway (Fig. 4A), we hypothesized that it could also play a role in the emergence of evaders. To investigate this hypothesis, we first measured intracellular ATP levels in Δ*purD* both in rich medium and after incubation in HIP. As expected, in both media, the ATP concentration measured for Δ*purD* was 5-fold lower than that measured for the parental strain (Fig. 4B).

**Fig 4.**
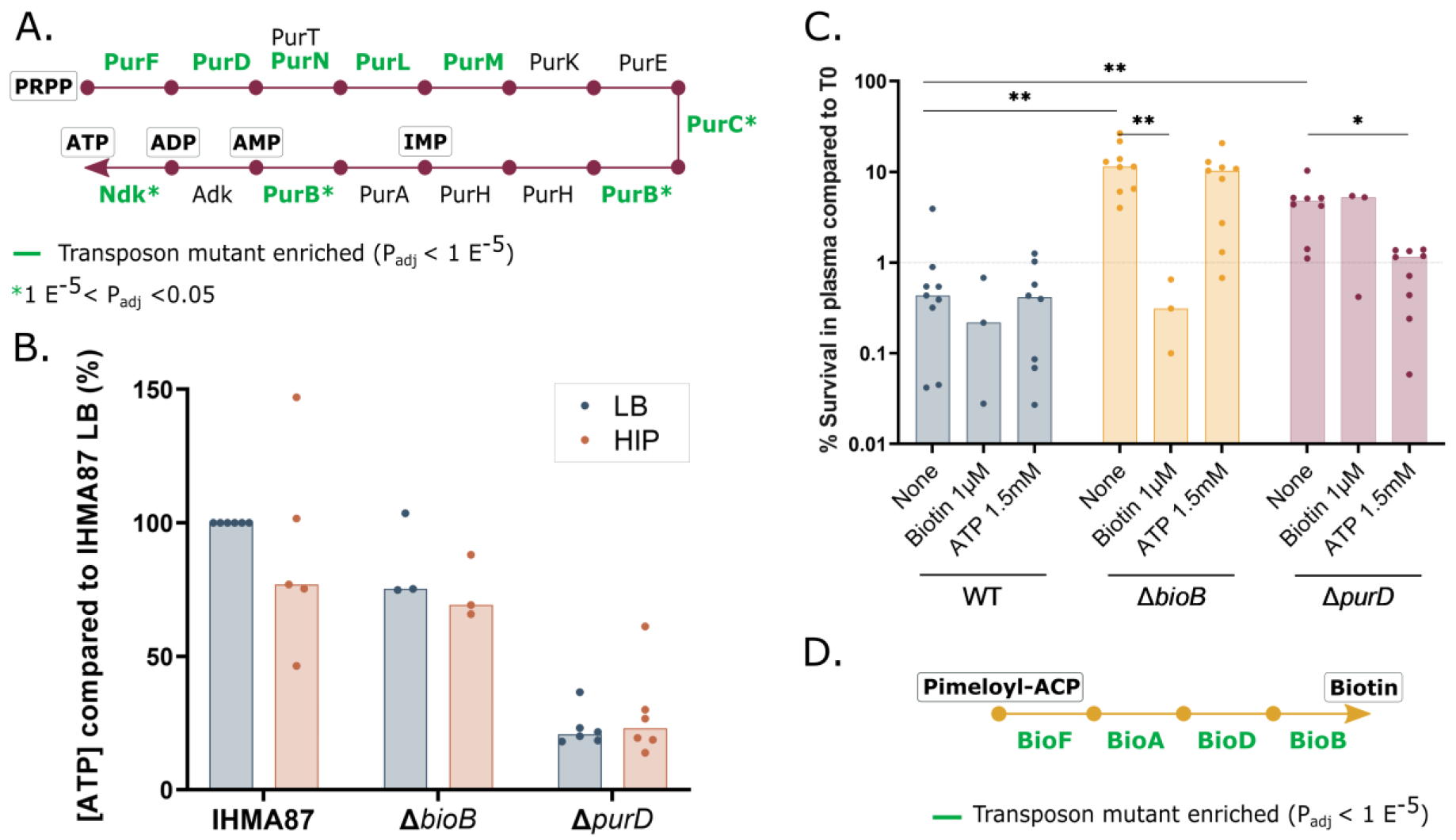
ATP and biotin influence bacterial resistance to plasma. **A.** Schematic view of the purine pathway (adapted from KEGG database (Kanehisa et al., 2008)). Significantly enriched insertions in corresponding genes of the Tn-seq screen are indicated**. B.** Measurement of intra-bacterial ATP levels in LB and after 2h-incubation in HIP normalized to the CFU counts. **C.** Trans-complementation of Δ*bioB* and Δ*purD* phenotype by exogenous biotin or ATP, respectively. Biotin (1 μM) and ATP (1.5 mM) were added at the beginning of the incubation of bacteria in plasma. The survival was estimated by CFU counting. **D.** The biotin biosynthetic pathway. Significantly enriched hits in Tn-seq are highlighted in green.

To confirm that the low ATP concentration was indeed responsible for higher resilience to human plasma, we modulated ATP levels by supplementing the plasma with exogenous ATP during the plasma killing assay (Fig. 4C). The addition of ATP (1.5 mM) when bacteria were exposed to plasma restored a wild-type-like sensitivity, further demonstrating the ATP-dependent phenotype of *pur* mutants.

No apparent links between the biotin biosynthesis pathway (Fig. 4D) and ATP synthesis exist. Nevertheless, as biotin is involved in many biological processes, we tested whether the Δ*bioB* mutant could be rescued by ATP. As shown in Figure 4C, biotin but not ATP restored a wild-type-like sensitivity to Δ*bioB*, suggesting a distinct molecular mechanism for evader formation in this mutant.

### Formation of energy-storage polyP granules in response to plasma

We then investigated potential physiological and/or morphological changes that could explain the increased survival of the Δ*purD* mutant. Transmission electron microscopy (TEM) images were acquired for the different strains grown in LB medium or after plasma challenge (Fig. 5A). When grown in LB, the parental strain and the Δ*bioB* and Δ*purD* mutants were of similar size, shape and apparent envelope thickness. Bacteria incubated in human plasma systematically presented electron-dense granules of variable sizes (Fig. 5A). The Δ*purD* mutant cells occasionally harbored small electron-dense granules even in LB (Fig. 5A, blue arrows). The size and number of the electron dense granules increased upon incubation of Δ*purD* in human plasma (Fig. 5A, red arrows). In parental and Δ*bioB* bacteria, granules only appeared after incubation in plasma. This result lends support to the conclusion that increased survival of Δ*bioB* in plasma is governed by a distinct mechanism to survival of the Δ*purD* strain (Fig. S2).

**Figure 5.**
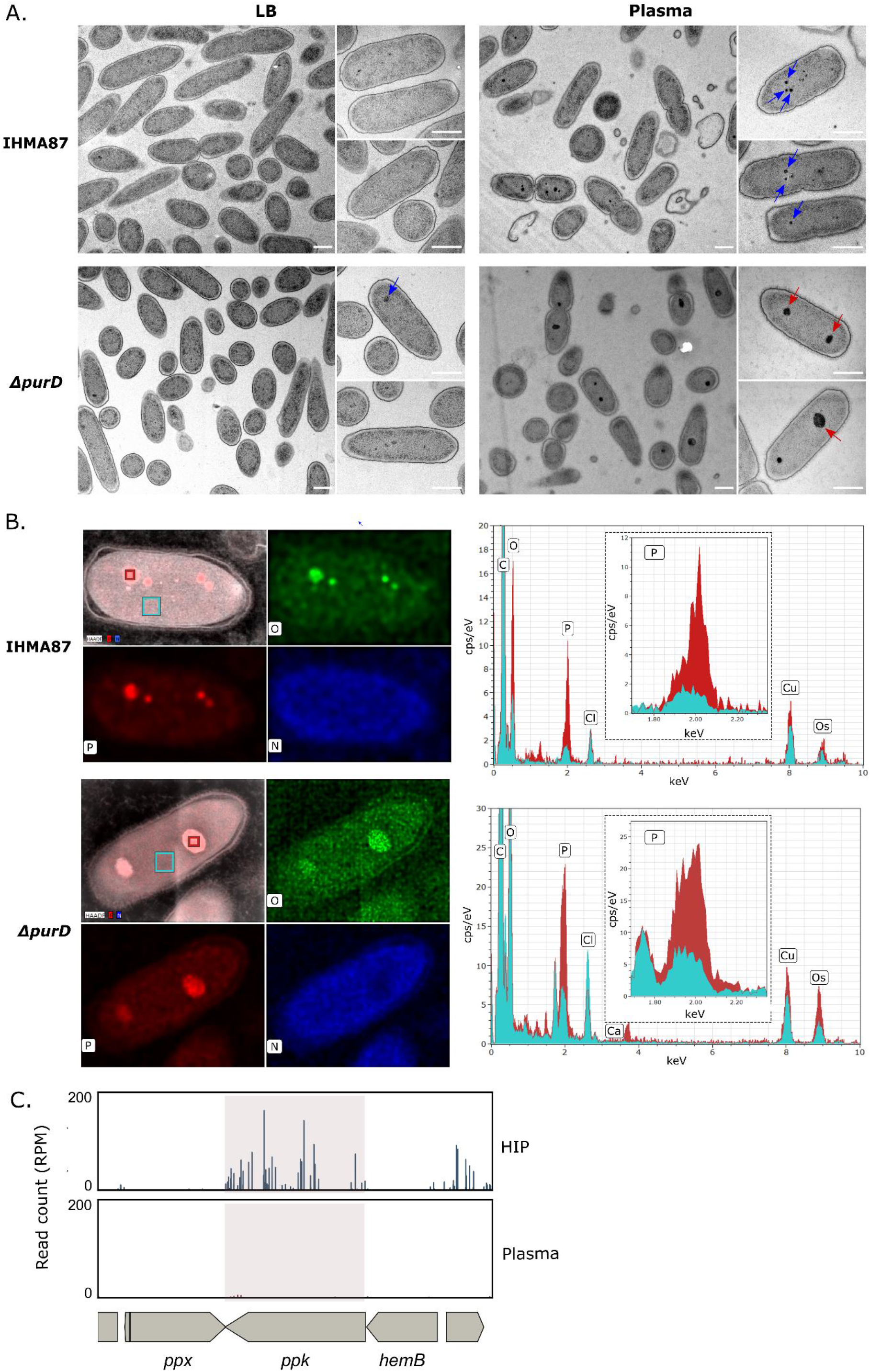
Evidence for the formation of polyP granules. **A.** Transmission electron microscopy images of indicated strains after growth in LB (left) or 1h-incubation in human plasma (right). Red and blue arrow show big- and small-sized granules respectively. Scale bar = 500 nm. **B.** EDX elemental composition maps of oxygen (green), phosphorus (red) and nitrogen (blue) of IHMA87 wild-type and Δ*purD* in plasma (left). EDX spectra (right) of highlighted locations (red = granules, cyan = cytoplasm), major peaks are assigned. Data show elevated levels of phosphorus and oxygen in the granule, indicating that the identified objects are polyP. **C.** Zoom in on Tn-seq profiles of *ppk1* gene with 2000 bp upstream and downstream, showing normalized numbers of reads in input (HIP) and output (plasma) samples.

Literature mining indicated that those electron-dense structures could be polyphosphate (polyP) granules. To cope with nutritional and energy stress, *P. aeruginosa* produces long polyP chains from both GTP and ATP. PolyP are then packed into granules to serve as energy and phosphate storage (Roewe et al, 2020; Racki et al., 2017). The granules detected here were confirmed to be phosphorus- and oxygen-rich structures by energy-dispersive X-ray spectroscopy (EDX) analysis (Fig. 5B). The formation of polyP granules upon incubation in plasma was also detected in the two BSI isolates, PaG1 and PaG7 (Fig. S2). As polyP formation could be part of the response to the active complement system or to the nutritive conditions of the plasma itself, we acquired images of the parental strain IHMA87 in heat-inactivated plasma (HIP). Granules were readily observed in bacteria incubated in HIP (Fig. S3).

PolyP are synthesized by polyphosphate kinases (Ppk), notably Ppk1 being the responsible for polyP synthesis in *P. aeruginosa* (Fraley et al., 2007). We hypothesized that the polyP could help *P. aeruginosa* to cope with plasma-related stress, and therefore we anticipate a deleterious effect of Ppk1 inactivation on IHMA87 survival in plasma. We then interrogated the Tn-seq data to evaluate the relevance of polyP in evaders’ formation. Indeed, sequencing profiles highlighted that transposon insertions in all of the three Ppk-encoding genes are deleterious in plasma (Log_2_(FC)= −2.33, Fig. 5C, Table 1S).

Altogether, incubation of *P. aeruginosa* in human plasma triggers the formation of dense polyP granules, even in the absence of complement system activity. In the plasma-tolerant strain, Δ*purD*, the granules observed were bigger and more numerous than in the parental strain. In addition, polyP biosynthetic genes are essential in plasma. Thus, the adaptation of bacteria to plasma through formation of energy storage granules may favor the selection of evaders that resist MAC killing.

### Specificity of mutant resistance toward MAC-induced killing

To examine the possible mechanisms deployed by selected mutants to resist the bactericidal activity of the MAC in plasma, we set up experiments to monitor several steps in the complement cascade. Complement activation by bacteria – both the classical and the alternative pathways – was examined by measuring the residual complement activity toward sensitized erythrocytes – sheep or rabbit, respectively – as described in (Dumestre-Pérard et al., 2008). After incubation with the different mutants, all plasma samples had the same residual complement capacity as plasma incubated with the parental strain (Fig. 6A-B), suggesting that the mutants did not alter complement activation.

**Fig 6.**
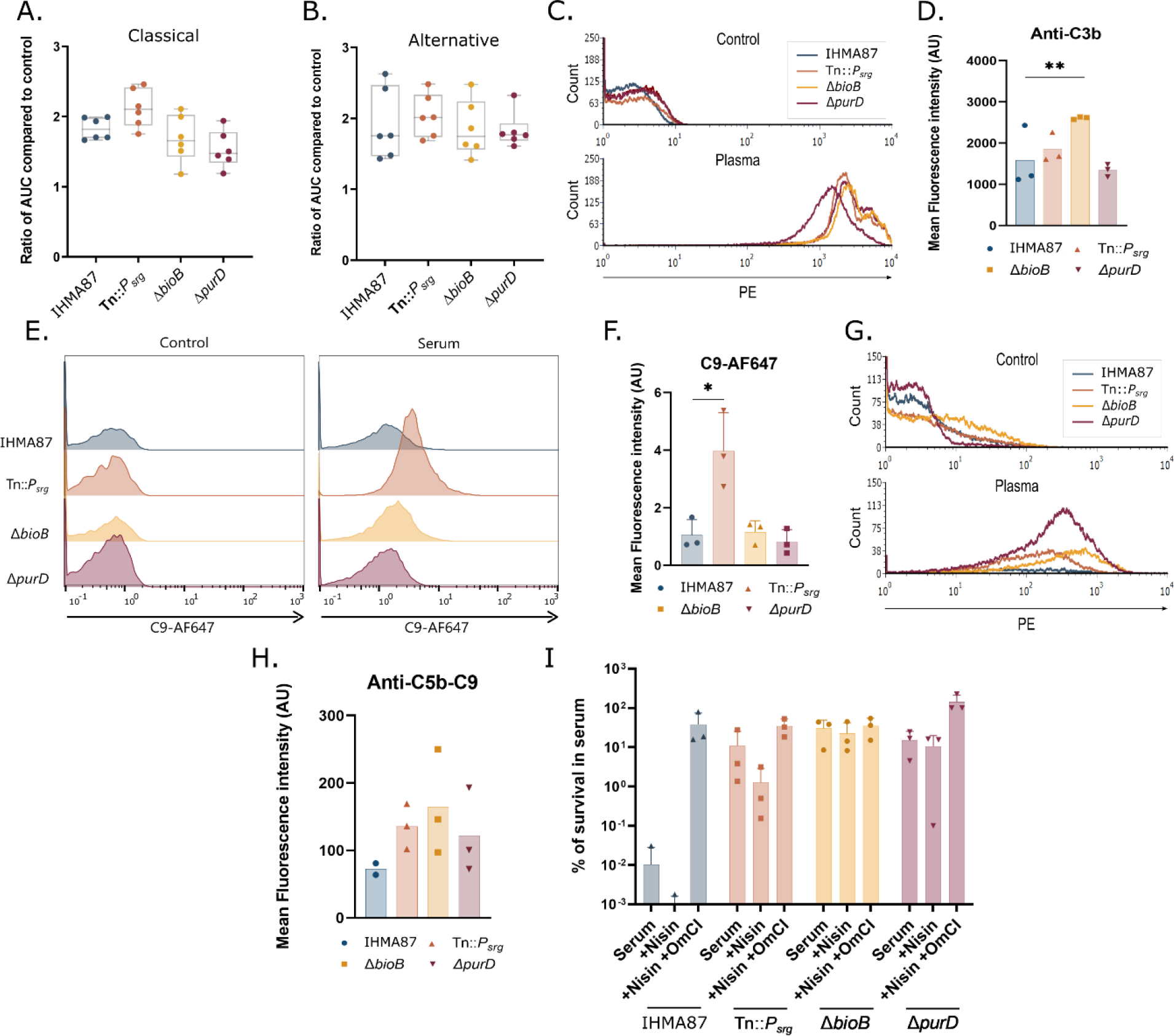
Mutants’ response to MAC-dependent killing. Residual complement activity of the classical (**A.**) and alternative (**B.**) complement pathways was measured after bacterial challenge in plasma, using lysis kinetics of erythrocytes, as described in Materials and Methods. Area under the curve (AUC) values were determined for each series and expressed relative to the 90% plasma pool control. n=3. **C.** C3b deposition on bacterial surface after incubation for 15 min in 0% (Control) or 90% plasma. Deposition was determined by FACS using the C3b specific PE-labeled antibody. **D.** Mean fluorescence intensities of the plasma condition presented in **C.** n=3. \*\**p*-value < 0.001. **E.** C9-AF647 insertion into bacterial membrane after 30 min in 0% (Control) or 3% pooled human serum and **F.** mean fluorescence intensities (\**p*-value < 0.05). **G.** C5b-9 deposition on bacterial surface after 15 min incubation in 0% (Control) and 90% plasma. Deposition was determined by FACS; mean fluorescence intensities are plotted (**H.**). n=3. **I.** Bacterial survival in pooled human serum or in the presence of nisin with or without the complement inhibitor OmCI.

To investigate whether the mutants prevent the formation of C3b and MAC (C5b-9) complexes, we first assessed deposits of CS components on the bacterial surface by flow cytometry (Fig. 6C-H). As shown in Figure 6C-D, increased C3b deposition was observed at 15 min for the Δ*bioB* mutant. When bacteria were exposed to pooled human serum supplemented with fluorescently-labeled C9 (Figure 6E-F), similar levels of C9 deposition were observed for most strains, with an apparent increase for the Tn∷*P_srg_* mutant.

Finally, we used flow cytometry with an anti-C5b-9 antibody to investigate MAC polymerization (Fig. 6G-H). No significant variation in the amount of C5b-9 complex was detected between the different mutants suggesting that, overall, MAC coating was unaltered in the mutants.

We therefore assessed the functionality of the MAC pores inserted in the outer membrane by comparing survival of the different bacterial strains in serum in the presence of nisin. The antimicrobial molecule nisin normally has no effect on Gram-negative bacteria with an intact outer membrane (Heesterbeek et al., 2019). However, MAC-induced membrane damage allows the compound to enter the bacterial cells, resulting in killing. As previously shown for plasma, wild-type *P. aeruginosa* is highly sensitive to killing in serum, with very efficient bacterial elimination (Fig. 6I). In contrast, increased survival was recorded for all mutants. Interestingly, the presence of nisin led to the death of most of the evader population. Similarly, numbers of the Tn∷*P_srg_* mutant were reduced in the presence of nisin, indicating that MAC-induced outer membrane damage does occur in this strain. These observations suggest that outer membrane damage by the MAC itself was not sufficient to kill this mutant or the evader population. In contrast, survival of the Δ*bioB* and Δ*purD* strains was unaltered by the addition of nisin to serum. Survival of all strains could be rescued by addition of C5 inhibitor OmCI, confirming that the killing is complement-mediated. Taken together, these data suggest that while the Δ*bioB*, Δ*purD*, and Tn∷*P_srg_* mutants survive better than the wild-type strain in serum, the Tn∷*P_srg_* mutant interferes at a different stage of MAC-dependent killing than the Δ*bioB* and Δ*purD* mutants.

## Discussion

The aim of this study was to elucidate the mechanisms deployed by *P. aeruginosa* to persist in human plasma. *P. aeruginosa* strains display highly variable survival in human plasma, ranging from fully resistant to sensitive. The majority of strains that are sensitive at the whole population level form a persistent sub-population with as yet uncharacterized features (Pont et al., 2020). Through this study, we identified ATP, biotin, and a small periplasmic protein – SrgA – as novel molecular determinants of *P. aeruginosa* resilience to the human CS. Other groups have performed genome-wide screens in complement-resistant strains of a number of pathogens in human serum (McCarthy et al., 2018; Phan et al., 2013; Poulsen et al., 2019; Sanchez-Larrayoz et al., 2017; Short et al., 2020). We found several of the factors identified, such as long OSA and exopolysaccharides, to also be important for the survival of *P. aeruginosa* in human plasma. Similarly, in agreement with genetic screens that identified bacterial persistence determinants following antibiotic treatment (Cameron et al., 2018; Molina-Quiroz et al., 2016), our results suggest that *P. aeruginosa* persistence in plasma is a multifactorial and probably stochastic phenomenon.

The mutant with reduced intracellular ATP levels (due to a mutation in the purine pathway) showed a high plasma-tolerance profile, and MAC-sensitivity could be restored by the external addition of ATP. In the context of higher tolerance/persistence to antibiotics, it has been proposed that the ATP-depleted conditions could decrease overall protein synthesis, and thus availability of antibiotic targets (Shan et al., 2017). Similarly, slow growth *per se*, whatever the restrictions causing it, can trigger a persistent state (Pontes and Groisman, 2019). Although incapable of growth, Δ*bioB* and Δ*purD* mutants survived and persisted in HIP for more than 24 hours. Admittedly, mutations in the purine and biotin pathways are unlikely to occur *in vivo*, but bacteria could use regulatory mechanisms to modulate purine and biotin synthesis, to enter into an antibiotic- or complement system-tolerant state. Furthermore, concentrations of ATP and biotin in the host may vary between individuals and health conditions. Moreover, some authors have suggested that the biotin biosynthesis pathway could be an alternative target for antibacterial agents (Carfrae et al., 2020). The biotin biosynthetic pathway and enhanced resistance to microbicidal activity of the CS revealed here suggest that the role of biotin in BSI should be carefully assessed before launching into any major developments.

The number of plasma evaders present following incubation in plasma varies between experiments, strains and the plasma used ((Pont et al., 2020) and this work). Therefore, it is possible that the initial incubation which is considered poorly nutritive and stressful for bacteria may stochastically trigger the emergence of evaders. Indeed, upon incubation in active or inactivated plasma, wild-type IHMA87 bacteria and BSI isolates form polyP granules that are energy storage structures (Roewe et al, 2020; Racki et al., 2017), and Δ*purD* mutant with low ATP levels displayed even bigger granules. Moreover, Ppk-encoding enzymes responsible for polyP synthesis are found critical for evaders formation, clearly suggesting that bacteria sense and respond to the stressful plasma environment. Upon incubation in human serum, bacteria upregulate hundreds of genes, the identity of which depends on the species. Interestingly, in plasma-resistant *P. aeruginosa* strain PAO1, exposure to human blood and serum triggers upregulation of quorum-sensing genes partly through the general virulence regulator Vfr (Beasley et al., 2020; Kruczek et al., 2014, 2016). However, the serum factors triggering expression of these genes and the downstream effectors in bacteria remain unknown.

In our search to identify the mechanisms used by the mutants to escape complement-mediated killing, we found that Tn∷*P_srg_*, Δ*bioB*, and Δ*purD* mutants do not impede complement activation. Rather, they probably modulate the function of MAC itself. MAC is an 18-unit oligomeric multiprotein pore formed by sequential addition of C5b, C6, C7, C8, and finally C9, to create a ring in the bacterial membrane (Bayly-Jones et al., 2017). Although the composition of MAC has been known for a number of years, its contribution to bacterial cell death and the mechanism causing bacterial death following MAC insertion remain debated (Doorduijn et al., 2019). Recent reports suggest that complement-resistant bacteria can specifically block C9 polymerization (Doorduijn et al., 2021). SrgA – 7.1-kDa after cleavage of its signal peptide – may directly interfere with MAC assembly. Alternatively, SrgA may act on peptidoglycan or other components of the Gram-negative envelope to impede MAC function. Despite several attempts, we have so far been unable to detect SrgA using protein tags in *P. aeruginosa*, but preliminary visualization of the protein in *E. coli* confirms that the *srgA* gene product is of a peptide nature. In contrast, the nature of the *srgB* and *srgC* gene products is completely speculative.

The role played by ATP in MAC activity is intriguing. No data are available on the energy needs for MAC assembly, but we can speculate that energy may be required for one of the steps during pore formation. Interestingly, although the majority of evaders were sensitive to the outer-membrane-impermeable polycyclic antibacterial peptide nisin (Heesterbeek et al., 2019), the mutants displayed resistance. Thus, in these mutants, the MAC pore was insufficient to allow nisin uptake through the permeabilized outer membrane. This difference in sensitivity could be due to defective pore insertion or to more resistant inner membranes. In evaders, in contrast, the outer membrane seems to have been effectively damaged by the MAC, suggesting that these bacteria have a more resistant envelope or can repair envelope damage more rapidly. Further characterization of evader biology and comparison to the bulk population will be needed to identify and further explore the molecular mechanisms leading to the emergence of evaders.

Given the genetic and phenotypic diversity of *P. aeruginosa* strains (Freschi et al., 2019; Hilker et al., 2015), and their variable levels of survival in human plasma (Pont et al., 2020), we expect that both common and strain-specific determinants counteracting MAC-dependent killing will be discovered. Only by screening several representative bacterial strains in parallel can we hope to obtain an overall picture of the *P. aeruginosa* plasma and blood resistome/persistome.

In an era where bacterial resistance to antibiotics has become a global health concern, it is highly relevant to explore any alternative means to eliminate pathogens. Harnessing the CS or MAC may be one possibility. For example, strategies to enhance MAC formation could be combined with classical antibiotics, phages, or antibodies (Abd El-Aziz et al., 2019; Cruz et al., 2019; Gulati et al., 2019). However, as yet, our knowledge of how various pathogens escape complement-mediated killing remains very limited. Better understanding of the molecular mechanisms involved in pathogen-complement interplay should help to design strategies to combat multi-drug resistant bacteria.

## Materials and methods

### Bacterial strains and genetic manipulations

Bacterial strains, plasmids, and primers are listed in Tables 3S and 4S. *E. coli* and *P. aeruginosa* were grown in LB at 37 °C with shaking (300 rpm). *P. aeruginosa* was selected on LB plates containing 25 μg/mL irgasan. Antibiotic concentrations were as follows: 75 μg/mL gentamicin, 75 μg/mL tetracycline, and 300 μg/mL carbenicillin for *P. aeruginosa*; and 50 μg/mL gentamicin, 10 μg/mL tetracycline, and 100 μg/mL ampicillin for *E. coli*. To create deletion mutants, upstream and downstream flanking regions (approximately 500 bp) were amplified from genomic DNA by PCR using appropriate primer pairs (sF1/sR1, sF2/sR2). Overlapping fragments were cloned into *Sma*I-digested pEXG2, pEX100T, or pEX18Tc by Sequence- and Ligation-Independent Cloning (SLIC, (Li and Elledge, 2007)). Plasmids were used to transform *E. coli* Top10 competent cells, and their sequences were verified (Eurofins). Allelic exchange vectors derived from pEXG2, pEX100T, or pEX18Tc were introduced into *P. aeruginosa* by triparental mating, using pRK600 as a helper plasmid. Merodiploids, resulting from homologous recombination, were selected on LB agar plates containing irgasan and the appropriate antibiotic. Single merodiploid colonies were streaked on NaCl-free LB plates with 10% sucrose (w/v) to select for plasmid loss. Resulting sucrose-resistant clones were screened for antibiotic sensitivity and gene deletion by PCR using appropriate primer pairs (F0/R0).

### Generation of the IHMA87 transposon library

The IHMA87 transposon mutant library was constructed essentially as previously described (Skurnik et al., 2013; Trouillon et al., 2020). Briefly, IHMA87, grown overnight at 42 °C under shaking, was mixed with two *E. coli* strains carrying pRK2013 (Figurski and Helinski, 1979) and pBTK24. The conjugation mixtures – containing 100 μL of each bacterial culture adjusted to an optical density at 600 nm (OD_600_) of 1 – were centrifuged, washed with LB, deposited on pre-warmed LB plates and incubated at 37 °C for 5 h. Approximately 100 conjugation mixtures yielded a library of > 300,000 mutants. Bacteria were scraped off into liquid LB, and aliquots were plated on LB plates containing irgasan (25 μg/mL) and gentamicin (75 μg/mL). Following growth at 37 °C, colonies were collected directly into LB containing 20% glycerol, aliquoted and stored at −80 °C.

### Preparation of pooled plasma

Pooled plasma was prepared from heparinized human healthy donor blood provided by the French National Blood Institute (EFS, Grenoble, France). Fresh blood was centrifuged for 10 min at 1,000 × g at room temperature. Supernatants from ten distinct donors were pooled, filtered through a 0.45-μm membrane and aliquoted prior to storage at −80 °C until needed. The same procedure was applied to citrated blood from 30 healthy donors. Before use, pooled plasma aliquots were thawed on ice, centrifuged for 10 min at 10,000 rpm and filtered through a 0.22-μm membrane. For the heat inactivated plasma (HIP) condition, plasma was thawed and heat inactivated for 30 min at 56 °C, centrifuged for 10 min at 10,000 rpm, and filtered through a 0.22-μm membrane.

### Plasma screening of *P. aeruginosa* strains

A vial containing the IHMA87 mutant library was thawed on ice, transferred to a flask containing 30 mL of LB and grown for 16 h under agitation at 27 °C, to limit bacterial growth. The next day, the bacterial suspension was diluted to OD_600nm_ ~0.1 and grown at 37 °C under agitation. Meanwhile, plasma and HIP were prepared. When cultures had reached an OD_600nm_ ~1, bacteria were harvested and resuspended in PBS supplemented with calcium and magnesium ions (Thermofisher Scientific) before exposing them to plasma, HIP, or LB at a concentration equivalent to 2.25×10^7^ CFU/mL (90% final plasma concentration). The precise initial CFU count for each experiment was determined by serial dilution. Samples were then incubated at 37 °C for 3 h on a rotating wheel. At the end of the challenge, an aliquot of each sample was transferred to LB and grown overnight; another aliquot was spread on LB agar plates to isolate individual mutants. Finally, bacterial survival rates were calculated based on the CFU counts for each sample. The same protocol was applied with the wild-type strain in parallel to compare survival rates. The next day, culture aliquots were harvested and stored for further mutant isolation. DNA was extracted from about 1×10^9^ bacteria for each sample.

### Illumina library construction, sequencing, and data analysis

#### Genomic DNA

Bacterial pellets from the screen in either HIP (Input) or plasma (Output) were used to prepare gDNA. A classical DNA-preparation protocol was applied, involving an SDS-NaCl lysis step (2% SDS, 0.15 M NaCl, 0.1 M EDTA pH8, 0.6 M Sodium perchlorate) followed by two successive phenol-chloroform extractions. The gDNA was precipitated with 100% cold ethanol and diluted in 200 μL TE buffer (10 mM Tris-HCl, 1 mM EDTA, pH8), producing a final concentration of 50-100 ng/μL. DNA was mechanically sheared at 4 °C with a Qsonica sonicator (Q700) in 15-s pulses for a total of 20 min. DNA fragments were then concentrated on columns (Monarch PCR & DNA Cleanup Kit, NEB). The size of the DNA fragments generated (150-400 bp) was verified on 2% agarose gel.

#### Library construction

Fragmented gDNA (1 μg) was end-repaired using an End-repair module (NEB#E6050), and dA-tailed using Klenow (NEB#6053). Short and Long adaptors (Table 1S) were annealed in 10 mM MgCl_2_ in a thermocycler programmed to decrease the temperature by 1 °C/cycle between 95 °C and 20 °C. gDNA fragments were ligated with annealed adaptors overnight at 16 °C in the presence of T4 DNA ligase (NEB#M0202). Bands from 200 to 400 bp were selected by extraction from a 2% agarose gel using a Monarch DNA Gel Extraction Kit (NEB#T1020L). Purified adaptor-DNA fragments were amplified by Phusion polymerase (NEB#M0530) in combination with specific primers (PCR1 Tn-spe direct and PCR1 Adaptor comp.). A second amplification round was performed with a primer bearing a P5 Illumina sequence and P7 indexed Illumina primers (NEB#E7335S). Primer sequences are available on request. After each step, products were quantified using a Qubit dsDNA HS Assay kit (Q33230). The quality of the libraries was assessed on an Agilent Bioanalyzer 2100 using high-sensitivity DNA chips (Ref #5067-4626). Constructs were sequenced on an Illumina NextSeq High (I2BC, Saclay Paris).

#### Data analysis

Sequencing reads were trimmed and aligned with the IHMA87 genome using Bowtie2 (Langmead and Salzberg, 2012). Htseq-count (Anders et al., 2015) was then used to determine read-counts for each feature. DESeq2 (Love et al., 2014) was applied to determine differential representation of insertion mutants between the plasma and HIP samples. To analyze insertions in intergenic regions, an annotation file was generated where intergenic regions were attributed to the upstream gene regardless of its orientation. The analysis protocol described above, using Htseq-count and DESeq2, was applied. Tn-seq was performed on biological duplicates.

### Plasma killing assay

Killing assays were performed as described (Pont et al., 2020). Unless stated otherwise, bacterial pre-cultures were diluted in LB to OD_600nm_ = 0.1 and grown at 37 °C under agitation until OD_600nm_ ~ 1. Bacteria were resuspended in PBS +/+ (Thermofisher Scientific) and incubated in plasma at a concentration of 2.25×10^7^ CFU/mL (90% final plasma concentration). The precise initial CFU count used in each experiment was determined by serial dilution. Bacterial survival rates were calculated based on CFU counts at the indicated time(s) and compared to the initial count. ATP, biotin, and purine were added to the plasma before adding bacteria.

### Transcriptomics

#### RNA extraction

Bacterial cultures were diluted to an OD_600nm_ of 0.1 and grown at 37 °C under agitation until they reached an OD_600nm_ of 1. RNA was isolated using the hot phenol-chloroform method. Briefly, bacteria were lysed in a hot (60 °C) phenol solution (2 mM EDTA, pH8, 1% SDS, 40 mM sodium acetate in acid phenol (Invitrogen #15594-047)). RNA was then isolated by successive phenol-chloroform extractions, followed by a cold chloroform extraction, and ethanol precipitation. Any residual genomic DNA was eliminated by Turbo DNase (Invitrogen #AM1907) treatment. Quantity and quality of total RNA were assessed using an Agilent Bioanalyzer.

#### Construction of RNA-seq libraries

Following RNA extraction, ribosomal RNAs (rRNAs) were depleted using a ribominus kit (Invitrogen) according to the manufacturer’s instructions. Depletion quality was verified on an Agilent Bioanalyzer using RNA pico chips before concentrating RNA by ethanol precipitation.

The NEBNext Ultra II directional RNA library prep kit for Illumina (NEB) was used to build cDNA libraries in preparation for sequencing, starting with 50 ng of rRNA-depleted RNA. The manufacturer’s instructions were followed. The quality and concentration of cDNA libraries were assessed using Agilent Bioanalyzer High-sensitivity DNA chips.

#### Sequencing and data analysis

DNA was sequenced at the Institute for Integrative Biology of the Cell (I2BC http://www.i2bc.paris-saclay.fr) high-throughput sequencing facility using an Illumina NextSeq500 instrument. More than 12 million 75-bp single-end reads were obtained per sample. Processed reads were mapped onto the IHMA87 genome (available on NCBI CP041354 and CP041355 accession numbers, for the chromosome and plasmid sequences, respectively (Trouillon et al., 2020)) using Bowtie2 (Langmead and Salzberg, 2012) and the readcount per feature was calculated using Ht-seq count (Anders et al., 2015). Finally, DESeq2 (Love et al., 2014) was applied to determine differential gene expression between the Tn∷*P_srg_* and wild-type strains. RNA-seq experiments were performed on biological duplicates.

### Alginate assay

Equivalent amounts of OD_600nm_-adjusted bacterial cultures were spread on *Pseudomonas* isolation agar (PIA) plates and incubated for 24 h at 37 °C. Bacterial lawns were flushed with saline solution (0.9% NaCl) and scraped off the plates. Bacteria were pelleted, and supernatant was stored in a separate tube. The bacterial pellet was resuspended in LB and incubated on a rotating wheel for 1-2 h to disrupt bacterial aggregates. The titer of the suspension was determined by serial dilution and CFU counting. Alginate was assayed as described in (Knutson and Jeanes, 1968; May and Chakrabarty, 1994). Briefly, 600 μL of ice-cold borate working solution (borate stock solution (4 M BO_3_^3−^) diluted to a final concentration of 100 mM in H2S04) was added to 70 μL of the bacterial supernatant, saline solution for a blank, and to commercial sodium alginate solution to produce a standard curve. On ice, 20 μL of carbazole solution (0.1% carbazole (Sigma #442506) in 100% ethanol) was added, and samples were incubated at 55 °C for 30 min after mixing. Finally, absorbance was measured at 530 nm.

### Determination of intracellular bacterial ATP concentration

To measure ATP in bacteria grown in LB, about 1 × 10^8^ bacteria were pelleted and resuspended in PBS. HIP was inoculated with a final concentration of 2.025 × 10^7^ bacteria/mL in PBS (90% final HIP concentration). Samples were incubated for 2 h at 37 °C on a rotating wheel, centrifuged and resuspended in PBS. Bacterial aggregates were gently dissociated by sonication using a Qsonica sonicator (Q700) for 45 sec at 10% intensity in 5-s-ON-5-s-OFF intervals. For the two conditions, 10-fold serial dilutions were prepared in 96-well plates, and precise bacterial counts were determined based on CFUs. The ATP concentration was determined using the BacTiter-Glo kit (Promega, #G8230). Briefly, equal volumes of bacterial suspension and BacTiter-Glo reagent were mixed in white 96-well Lumitrac plates (Bio-Rad #655075) and incubated for 5 min under shaking. A standard curve (0.1 to 100 nM ATP) was established for each experiment. Bioluminescence was measured on a plate reader (Spark, TECAN). The ATP concentration was then expressed relative to the CFU count. Experiments were performed in more than three biological replicates.

### Residual complement activity

#### Sample preparation

Bacteria grown to 0D_600nm_ ~ 1 were resuspended in PBS +/+ (Thermofisher Scientific) and incubated in plasma at a concentration of 2.25×10^9^ CFU/mL (90% final plasma concentration), or PBS alone as control. Samples were incubated for 3 h at 37 °C on a rotating wheel to trigger complement activation by bacteria. The initial CFU count was determined by serial dilution. At the end of the incubation, samples were centrifuged and supernatant was filtered through a 0.22-μm membrane to remove any remaining bacteria. Samples were aliquoted and frozen at −80 °C until use. Residual complement activity in the bacterial/plasma supernatant was measured using a total hemolytic assay to study classical and alternative pathways, as described in (Dumestre-Pérard et al., 2008).

#### Classical complement hemolytic activity

Briefly, sheep erythrocytes (Orgentec) were sensitized using anti-erythrocyte antibodies (1/40 000; Hemolysin, Orgentec) for 15 min at 37 °C. Bacterial/plasma supernatant (25 μL) was mixed with 3 mL of sensitized erythrocytes (4×10^6^ cells/mL) diluted in DGVB^2+^ (2.5% glucose, 0.05% gelatin, 2.5 mM Veronal, 72.5 mM NaCl, 0.15 mM Ca^2+^, 0.5 mM Mg^2+^), and lysis kinetics were monitored based on OD_660nm_ using a spectrophotometer (Safas UVmc2). Area under the curve values were determined for each series and expressed relative to the 90% plasma pool control.

#### Alternative complement hemolytic activity

Briefly, rabbit erythrocytes (Ecole Nationale Vétérinaire de Toulouse) were washed three times using DGVB-Mg-EGTA (2.5% glucose, 0.05% gelatin, 2.5 mM Veronal, 72.5 mM NaCl, 5 mM Mg^2+^, 3 mM EGTA). Bacterial/plasma supernatant (150 μL) was added to 3 mL of a rabbit erythrocyte suspension (diluted in DGVB-Mg-EGTA at 1/250) at 37 °C, and lysis kinetics were monitored as described for classical complement activity.

### C3b and C5b-9 binding assays

Bacteria (4 × 10^7^) were incubated for 15 min in a 90% heparinized plasma pool at 37 °C. Incubation was stopped by diluting the bacterial suspension in ten volumes of PBS containing 20 mM EDTA (pH8). C3b binding was detected with PE-conjugated mouse monoclonal anti-C3b/iC3b (5 μg/mL, BioLegend #846103) and C5b-9 membrane deposition/insertion was detected with mouse monoclonal anti-C5b-9 antibody (8 μg/mL, Abcam #ab66768). Antibody binding was revealed with PE-conjugated goat anti-mouse antibodies (2.5 μg/mL, Abcam #ab97041). The negative control consisted of bacteria in 0% plasma (100% PBS). Samples were analyzed on a FACSCalibur (Becton Dickinson); data were acquired and analyzed using FCSExpress6 Software.

### C9 binding

*P. aeruginosa* strains expressing green fluorescent protein (GFP) under the control of the constitutive *P_X2_* promoter were grown overnight in LB supplemented tetracycline at 37 °C, with shaking. Bacteria were subcultured 1:30 in LB supplemented with the antibiotic and allowed to grow to a mid-log OD_600nm_ of 0.5 at 37 °C, with shaking. Bacteria were pelleted by centrifugation, washed, and resuspended at a final OD_600nm_ of 0.05 in RPMI + 0.05% human serum albumin (HSA) for the experiments. Bacteria were then exposed to pooled human serum supplemented with 100 nM C9-AF647 and incubated for 30 min at 37 °C, with shaking. To count CFU, bacteria were serially diluted in MilliQ water, spotted on LB plates, and incubated overnight at 37 °C. For flow cytometry, bacteria were diluted 10-fold in RPMI-HSA and analyzed by flow cytometry (MACSquant VYB; Miltenyi Biotech). Bacteria were gated on GFP-expression and subsequently analyzed for AF647 fluorescence. Flow cytometry data was analyzed using FlowJo V.10.

### Nisin assay

Bacteria were grown overnight in LB at 37 °C, with shaking, subcultured 1:30 in LB to a mid-log OD_600nm_ of 0.5 at 37 °C, with shaking, pelleted by centrifugation, washed, and resuspended at a final OD_600nm_ of 0.05 in RPMI + 0.05% HSA for the experiments. Bacteria were then exposed to human pooled plasma supplemented with buffer, 3 μg/mL nisin, or 3 μg/mL nisin and 10 μg/mL OmCI, and incubated for 3 h at 37 °C. Serial dilutions in MilliQ water were spotted on LB plates for incubation overnight at 37 °C to determine CFU the next day.

### Transmission electron microscopy

Bacterial pre-cultures were diluted to an OD_600nm_ of 0.1 in flasks containing 30 mL of LB and incubated at 37 °C, with shaking until OD_600nm_ of 1. Plasma was then inoculated with bacterial suspension to reach a bacterial concentration of 1.125 ×10^9^ bacteria/mL and incubated on a rotating wheel for 1 h at 37 °C. At the end of the challenge, bacteria were pelleted by centrifugation and resuspended in LB prior to fixation. In parallel, bacteria grown in LB to an OD_600nm_ of 1 were pelleted by centrifugation and resuspended in LB prior to fixation.

Samples were prepared as described in (Mohamed et al., 2021). Briefly, the cell pellet was spread on the 200-μm side of a 3-mm type A gold plate (Leica Microsystems) covered with the flat side of a 3-mm type B aluminum plate (Leica Microsystems) and vitrified by high-pressure freezing in a HPM100 system (Leica Microsystems). After freezing at −90 °C in acetone with 1% OsO_4_, samples were slowly warmed to −60 °C before storing for 12 h. Then, the temperature was increased to −30 °C and the samples were stored for a further 12 h. Samples were then warmed to 0 °C, incubated for 1 h, and cooled once again to −30 °C (AFS2; Leica Microsystems). Vitrified samples were rinsed four times in pure acetone before infiltration with progressively increasing concentrations of resin (Epoxy embedding medium, Sigma) in acetone while increasing the temperature to 20 °C. Pure resin was added at room temperature. After polymerization, thin (70-nm) sections were prepared using a UC7 ultramicrotome (Leica Microsystems) and collected on 100-mesh copper grids coated with Formvar carbon. The thin sections were post-stained for 5 min with 2% uranyl acetate, rinsed with water and incubated for 2 min with lead citrate. Samples were observed using a Tecnai G2 Spirit BioTwin microscope (FEI) operating at 120 kV with an Orius SC1000B CCD camera (Gatan).

### Granules analysis

For the chemical analysis of granules, 200 nm-thin sections, collected on carbon/formvar coated copper grids, were observed in scanning/transmission electron microscopy (STEM) mode and analyzed by STEM (HAADF) detector, on a Tecnai OSIRIS microscope (FEI) operated at 200 kV. The chemical composition of regions of interest was then analyzed by EDS using the ESPRIT 1.9 software (Bruker).

### Bioinformatics analysis and protein conservation

Protein sequence files from all *P. aeruginosa* strains with a closed assembled genome were retrieved from the Pseudomonas genome database, and *srg* conservation was investigated using the Blast Reciprocal Best Hits program (Cock et al., 2015), selecting for 90% coverage and at least 50% identity.

Signal peptides were predicted on the SignalP website (Almagro Armenteros et al., 2019).

### Statistical analysis

Sigmaplot software was used to perform statistical analysis. Multiple-group comparisons were performed by applying one-way ANOVA or Kruskal-Wallis tests, depending on whether data was normally distributed. Subsequently, a Student-Newman-Keuls pairwise comparison was applied. In some cases (as indicated), non-normally-distributed data were converted to normally-distributed datasets by log10-transformation. Figures were designed using GraphPad Prism9.

## Data availability

The Tn-seq and RNA-seq data generated in this work are available through the NCBI Gene Expression Omnibus (GEO) under super-series accession number GSE192831, or by using accession numbers GSE192769 and GSE192761, respectively.

## Acknowledgments

The work described in this paper was supported by grants from the French national agency for research (Agence Nationale de la Recherche; ANR-15-CE11-0018-01), the Laboratory of Excellence GRAL, funded through the University Grenoble Alpes graduate school (Écoles Universitaires de Recherche) CBH-EUR-GS (ANR-17-EURE-0003), the Fondation pour la Recherche Médicale (Team FRM 2017, DEQ20170336705) to I.A., the European Union’s Horizon 2020 research programs H2020-EU-ITN-EJD (CORVOS #860044 to F.M. and SHMR). The sequencing experiments benefited from the facilities and expertise of the high throughput sequencing core facility at I2BC (Centre de Recherche de Gif – http://www.i2bc.paris-saclay.fr/). The *P. aeruginosa* IHMA87 strain was obtained from the International Health Management Association, USA. This work availed of the platforms at the Grenoble Instruct-ERIC center (ISBG; UAR 3518 CNRS-CEA-UGA-EMBL) within the Grenoble Partnership for Structural Biology (PSB), supported by FRISBI (ANR-10-INBS-0005-02) and GRAL, funded through the University Grenoble Alpes graduate school (Ecoles Universitaires de Recherche) CBH-EUR-GS (ANR-17-EURE-0003). S.P, J.T and M.JM were recipients of Ph.D. fellowships from the French Ministry of Education and Research. S.S received the Master 2 GRAL fellowship. The IBS Electron Microscope facility is supported by the Auvergne Rhône-Alpes Region, the Fonds Feder, the Fondation pour la Recherche Médicale, and GIS-IBiSA. The authors thank Emily Rey for her help with mutant analyses. M.JM thanks Twitter followers for indicating literature on PolyP granules.

## Bibliography

Abd El-Aziz, A.M., Elgaml, A., and Ali, Y.M.(2019). Bacteriophage Therapy Increases Complement-Mediated Lysis of Bacteria and Enhances Bacterial Clearance After Acute Lung Infection With Multidrug-Resistant Pseudomonas aeruginosa. J. Infect. Dis. 219, 1439–1447. https://doi.org/10.1093/infdis/jiy678.

Almagro Armenteros, J. J., Tsirigos, K.D., Sønderby, C.K., Petersen, T.N., Winther, O., Brunak, S., von Heijne, G., and Nielsen, H.(2019). SignalP 5.0 improves signal peptide predictions using deep neural networks. Nat. Biotechnol. 37, 420–423. https://doi.org/10.1038/s41587-019-0036-z.

Anders, S., Pyl, P.T., and Huber, W.(2015). HTSeq—a Python framework to work with high-throughput sequencing data. Bioinformatics 31, 166–169. https://doi.org/10.1093/bioinformatics/btu638.

Andersen-Civil, A.I.S., Ahmed, S., Guerra, P.R., Andersen, T.E., Hounmanou, Y.M.G., Olsen, J.E., and Herrero-Fresno, A. (2018). The impact of inactivation of the purine biosynthesis genes, purN and purT, on growth and virulence in uropathogenic E. coli. Mol. Biol. Rep. 45, 2707–2716. https://doi.org/10.1007/s11033-018-4441-z.

Ayrapetyan, M., Williams, T.C., Baxter, R., and Oliver, J.D.(2015). Viable but Nonculturable and Persister Cells Coexist Stochastically and Are Induced by Human Serum. Infect. Immun. 83, 4194–4203. https://doi.org/10.1128/IAI.00404-15.

Balaban, N.Q., Helaine, S., Lewis, K., Ackermann, M., Aldridge, B., Andersson, D.I., Brynildsen, M.P., Bumann, D., Camilli, A., Collins, J.J., et al. (2019). Definitions and guidelines for research on antibiotic persistence (vol 17, pg 441, 2019). Nat. Rev. Microbiol. 17, 460–460. https://doi.org/10.1038/s41579-019-0207-4.

Bartell, J.A., Cameron, D.R., Mojsoska, B., Haagensen, J.A.J., Pressler, T., Sommer, L.M., Lewis, K., Molin, S., and Johansen, H.K.(2020). Bacterial persisters in long-term infection: Emergence and fitness in a complex host environment. PLoS Pathog. 16, e1009112. https://doi.org/10.1371/journal.ppat.1009112.

Bayly-Jones, C., Bubeck, D., and Dunstone, M.A.(2017). The mystery behind membrane insertion: a review of the complement membrane attack complex. Philos. Trans. R. Soc. B Biol. Sci. 372, 20160221. https://doi.org/10.1098/rstb.2016.0221.

Beasley, K.L., Cristy, S.A., Elmassry, M.M., Dzvova, N., Colmer-Hamood, J.A., and Hamood, A.N.(2020). During bacteremia, Pseudomonas aeruginosa PAO1 adapts by altering the expression of numerous virulence genes including those involved in quorum sensing. PLOS ONE 15, e0240351. https://doi.org/10.1371/journal.pone.0240351.

de Bentzmann, S., Giraud, C., Bernard, C.S., Calderon, V., Ewald, F., Plésiat, P., Nguyen, C., Grunwald, D., Attree, I., Jeannot, K., et al. (2012). Unique Biofilm Signature, Drug Susceptibility and Decreased Virulence in Drosophila through the Pseudomonas aeruginosa Two-Component System PprAB. PLoS Pathog. 8, e1003052. https://doi.org/10.1371/journal.ppat.1003052.

Blom, A.M., Rytkönen, A., Vasquez, P., Lindahl, G., Dahlbäck, B., and Jonsson, A.-B. (2001). A Novel Interaction Between Type IV Pili of Neisseria gonorrhoeae and the Human Complement Regulator C4b-Binding Protein. J. Immunol. 166, 6764–6770. https://doi.org/10.4049/jimmunol.166.11.6764.

Broder, U.N., Jaeger, T., and Jenal, U.(2017). LadS is a calcium-responsive kinase that induces acute-to-chronic virulence switch in Pseudomonas aeruginosa. Nat. Microbiol. 2, 16184. https://doi.org/10.1038/nmicrobiol.2016.184.

Cameron, D.R., Shan, Y., Zalis, E.A., Isabella, V., and Lewis, K.(2018). A Genetic Determinant of Persister Cell Formation in Bacterial Pathogens. J. Bacteriol. 200. https://doi.org/10.1128/JB.00303-18.

Carfrae, L.A., MacNair, C.R., Brown, C.M., Tsai, C.N., Weber, B.S., Zlitni, S., Rao, V.N., Chun, J., Junop, M.S., Coombes, B.K., et al. (2020). Mimicking the human environment in mice reveals that inhibiting biotin biosynthesis is effective against antibiotic-resistant pathogens. Nat. Microbiol. 5, 93–101. https://doi.org/10.1038/s41564-019-0595-2.

Chiang, S.L., Taylor, R.K., Koomey, M., and Mekalanos, J.J.(1995). Single amino acid substitutions in the N-terminus of Vibrio cholerae TcpA affect colonization, autoagglutination, and serum resistance. Mol. Microbiol. 17, 1133–1142. https://doi.org/10.1111/j.1365-2958.1995.mmi_17061133.x.

Cock, P.J.A., Chilton, J.M., Grüning, B., Johnson, J.E., and Soranzo, N.(2015). NCBI BLAST+ integrated into Galaxy. GigaScience 4, s13742-015-0080–0087. https://doi.org/10.1186/s13742-015-0080-7.

Conlon, B.P., Rowe, S.E., Gandt, A.B., Nuxoll, A.S., Donegan, N.P., Zalis, E.A., Clair, G., Adkins, J.N., Cheung, A.L., and Lewis, K.(2016). Persister formation in Staphylococcus aureus is associated with ATP depletion. Nat. Microbiol. 1, 1–7. https://doi.org/10.1038/nmicrobiol.2016.51.

Cruz, J.W., Damko, E., Modi, B., Tu, N., Meagher, K., Voronina, V., Gartner, H., Ehrlich, G., Rafique, A., Babb, R., et al. (2019). A novel bispecific antibody platform to direct complement activity for efficient lysis of target cells. Sci. Rep. 9, 12031. https://doi.org/10.1038/s41598-019-48461-1.

Damron, F.H., and Goldberg, J.B.(2012). Proteolytic regulation of alginate overproduction in Pseudomonas aeruginosa: Proteolytic regulation of alginate. Mol. Microbiol. 84, 595–607. https://doi.org/10.1111/j.1365-2958.2012.08049.x.

Doorduijn, D.J., Rooijakkers, S.H.M., and Heesterbeek, D.A.C.(2019). How the Membrane Attack Complex Damages the Bacterial Cell Envelope and Kills Gram-Negative Bacteria. BioEssays 41, 1900074. https://doi.org/10.1002/bies.201900074.

Doorduijn, D.J., Heesterbeek, D.A.C., Ruyken, M., Haas, C.J.C. de, Stapels, D.A.C., Aerts, P.C., Rooijakkers, S.H.M., and Bardoel, B.W.(2021). Polymerization of C9 enhances bacterial cell envelope damage and killing by membrane attack complex pores. PLOS Pathog. 17, e1010051. https://doi.org/10.1371/journal.ppat.1010051.

Dumestre-Pérard, C., Lamy, B., Aldebert, D., Lemaire-Vieille, C., Grillot, R., Brion J.-P. Gagnon, J., and Cesbron, J.-Y. (2008). *Aspergillus* Conidia Activate the Complement by the Mannan-Binding Lectin C2 Bypass Mechanism. J. Immunol. 181, 7100–7105. https://doi.org/10.4049/jimmunol.181.10.7100.

Fauvart, M., De Groote, V.N., and Michiels, J.(2011). Role of persister cells in chronic infections: clinical relevance and perspectives on anti-persister therapies. J. Med. Microbiol. 60, 699–709. https://doi.org/10.1099/jmm.0.030932-0.

Figurski, D.H., and Helinski, D.R.(1979). Replication of an origin-containing derivative of plasmid RK2 dependent on a plasmid function provided in trans. Proc. Natl. Acad. Sci. U. S. A. 76, 1648–1652. .

Fraley, C.D., Rashid, M.H., Lee, S.S.K., Gottschalk, R., Harrison, J., Wood, P.J., Brown, M.R.W., and Kornberg, A.(2007). A polyphosphate kinase 1 (*ppk1*) mutant of *Pseudomonas aeruginosa* exhibits multiple ultrastructural and functional defects. Proc. Natl. Acad. Sci. 104, 3526–3531. https://doi.org/10.1073/pnas.0609733104.

Freschi, L., Vincent, A.T., Jeukens, J., Emond-Rheault, J.-G., Kukavica-Ibrulj, I., Dupont M.-J. Charette, S.J., Boyle, B., and Levesque, R.C.(2019). The *Pseudomonas aeruginosa* Pan-Genome Provides New Insights on Its Population Structure, Horizontal Gene Transfer, and Pathogenicity. Genome Biol. Evol. 11, 109–120. https://doi.org/10.1093/gbe/evy259.

Gaca, A.O., Colomer-Winter, C., and Lemos, J.A.(2015). Many Means to a Common End: the Intricacies of (p)ppGpp Metabolism and Its Control of Bacterial Homeostasis. J. Bacteriol. 197, 1146–1156. https://doi.org/10.1128/JB.02577-14.

Giraud, C., Bernard, C.S., Calderon, V., Yang, L., Filloux, A., Molin, S., Fichant, G., Bordi, C., and de Bentzmann, S. (2011). The PprA-PprB two-component system activates CupE, the first non-archetypal Pseudomonas aeruginosa chaperone-usher pathway system assembling fimbriae: P. aeruginosa CupE fimbriae. Environ. Microbiol. 13, 666–683. https://doi.org/10.1111/j.1462-2920.2010.02372.x.

Goldberg, J.B., and Pler, G.B.(1996). Pseudomonas aeruginosa lipopolysaccharides and pathogenesis. Trends Microbiol. 4, 490–494. https://doi.org/10.1016/s0966-842x(97)82911-3.

Gulati, S., Beurskens, F.J., de Kreuk, B.-J., Roza, M., Zheng, B., DeOliveira, R.B., Shaughnessy, J., Nowak, N.A., Taylor, R.P., Botto, M., et al. (2019). Complement alone drives efficacy of a chimeric antigonococcal monoclonal antibody. PLOS Biol. 17, e3000323. https://doi.org/10.1371/journal.pbio.3000323.

Harms, A., Maisonneuve, E., and Gerdes, K.(2016). Mechanisms of bacterial persistence during stress and antibiotic exposure. Science 354. https://doi.org/10.1126/science.aaf4268.

Heesterbeek, D. a. C., Martin, N.I., Velthuizen, A., Duijst, M., Ruyken, M., Wubbolts, R., Rooijakkers, S.H.M., and Bardoel, B.W.(2019). Complement-dependent outer membrane perturbation sensitizes Gram-negative bacteria to Gram-positive specific antibiotics. Sci. Rep. 9, 3074. https://doi.org/10.1038/s41598-019-38577-9.

Helaine, S., Cheverton, A.M., Watson, K.G., Faure, L.M., Matthews, S.A., and Holden, D.W.(2014). Internalization of Salmonella by macrophages induces formation of nonreplicating persisters. Science 343, 204–208. https://doi.org/10.1126/science.1244705.

Hilker, R., Munder, A., Klockgether, J., Losada, P.M., Chouvarine, P., Cramer, N., Davenport, C.F., Dethlefsen, S., Fischer, S., Peng, H., et al. (2015). Interclonal gradient of virulence in the Pseudomonas aeruginosa pangenome from disease and environment. Environ. Microbiol. 17, 29–46. https://doi.org/10.1111/1462-2920.12606.

Hill, P.W.S., Moldoveanu, A.L., Sargen, M., Ronneau, S., Glegola-Madejska, I., Beetham, C., Fisher, R.A., and Helaine, S.(2021). The vulnerable versatility of Salmonella antibiotic persisters during infection. Cell Host Microbe https://doi.org/10.1016/j.chom.2021.10.002.

Jones, C.J., and Wozniak, D.J.(2017). Psl Produced by Mucoid Pseudomonas aeruginosa Contributes to the Establishment of Biofilms and Immune Evasion. MBio 8, e00864-17, /mbio/8/3/e00864-17.atom. https://doi.org/10.1128/mBio.00864-17.

Kanehisa, M., Araki, M., Goto, S., Hattori, M., Hirakawa, M., Itoh, M., Katayama, T., Kawashima, S., Okuda, S., Tokimatsu, T., et al. (2008). KEGG for linking genomes to life and the environment. Nucleic Acids Res. 36, D480–D484. https://doi.org/10.1093/nar/gkm882.

Kang, C.-I., Kim, S.-H., Kim, H.-B., Park, S.-W., Choe, Y.-J., Oh, M., Kim, E.-C., and Choe, K.-W. (2003). Pseudomonas aeruginosa Bacteremia: Risk Factors for Mortality and Influence of Delayed Receipt of Effective Antimicrobial Therapy on Clinical Outcome. Clin. Infect. Dis. 37, 745–751. https://doi.org/10.1086/377200.

Keren, I., Kaldalu, N., Spoering, A., Wang, Y., and Lewis, K.(2004). Persister cells and tolerance to antimicrobials. FEMS Microbiol. Lett. 230, 13–18. https://doi.org/10.1016/S0378-1097(03)00856-5.

Kintz, E., Scarff, J.M., DiGiandomenico, A., and Goldberg, J.B.(2008). Lipopolysaccharide O-Antigen Chain Length Regulation in Pseudomonas aeruginosa Serogroup O11 Strain PA103. J. Bacteriol. 190, 2709–2716. https://doi.org/10.1128/JB.01646-07.

Knutson, C.A., and Jeanes, A.(1968). A new modification of the carbazole analysis: application to heteropolysaccharides. Anal Biochem 24, 470–481. https://doi.org/10.1016/0003-2697(68)90154-1.

Korch, S.B., Henderson, T.A., and Hill, T.M.(2003). Characterization of the hipA7 allele of Escherichia coli and evidence that high persistence is governed by (p)ppGpp synthesis. Mol. Microbiol. 50, 1199–1213. https://doi.org/10.1046/j.1365-2958.2003.03779.x.

Kruczek, C., Qaisar, U., Colmer-Hamood, J.A., and Hamood, A.N.(2014). Serum influences the expression of Pseudomonas aeruginosa quorum-sensing genes and QS-controlled virulence genes during early and late stages of growth. MicrobiologyOpen 3, 64–79. https://doi.org/10.1002/mbo3.147.

Kruczek, C., Kottapalli, K.R., Dissanaike, S., Dzvova, N., Griswold, J.A., Colmer-Hamood, J.A., and Hamood, A.N.(2016). Major Transcriptome Changes Accompany the Growth of Pseudomonas aeruginosa in Blood from Patients with Severe Thermal Injuries. PLOS ONE 11, e0149229. https://doi.org/10.1371/journal.pone.0149229.

Langmead, B., and Salzberg, S.L.(2012). Fast gapped-read alignment with Bowtie 2. Nat. Methods 9, 357–359. https://doi.org/10.1038/nmeth.1923.

Li, M.Z., and Elledge, S.J.(2007). Harnessing homologous recombination in vitro to generate recombinant DNA via SLIC. Nat. Methods 4, 251–256. https://doi.org/10.1038/nmeth1010.

Love, M.I., Huber, W., and Anders, S.(2014). Moderated estimation of fold change and dispersion for RNA-seq data with DESeq2. Genome Biol. 15, 550. https://doi.org/10.1186/s13059-014-0550-8.

Manuse, S., Shan, Y., Canas-Duarte, S.J., Bakshi, S., Sun, W.-S., Mori, H., Paulsson, J., and Lewis, K.(2021). Bacterial persisters are a stochastically formed subpopulation of low-energy cells. PLOS Biol. 19, e3001194. https://doi.org/10.1371/journal.pbio.3001194.

May, T.B., and Chakrabarty, A.M.(1994). [22] Isolation and assay of Pseudomonas aeruginosa alginate. In Methods in Enzymology, (Academic Press), pp. 295–304.

McCarthy, A.J., Stabler, R.A., and Taylor, P.W.(2018). Genome-Wide Identification by Transposon Insertion Sequencing of Escherichia coli K1 Genes Essential for In Vitro Growth, Gastrointestinal Colonizing Capacity, and Survival in Serum. J. Bacteriol. 200. https://doi.org/10.1128/JB.00698-17.

Mishra, M., Byrd, M.S., Sergeant, S., Azad, A.K., Parsek, M.R., McPhail, L., Schlesinger, L.S., and Wozniak, D.J.(2012). Pseudomonas aeruginosa Psl polysaccharide reduces neutrophil phagocytosis and the oxidative response by limiting complement-mediated opsonization. Cell. Microbiol. 14, 95–106. https://doi.org/10.1111/j.1462-5822.2011.01704.x.

Mohamed, A.M.T., Chan, H., Luhur, J., Bauda, E., Gallet, B., Morlot, C., Cole, L., Awad, M., Crawford, S., Lyras, D., et al. (2021). Chromosome Segregation and Peptidoglycan Remodeling Are Coordinated at a Highly Stabilized Septal Pore to Maintain Bacterial Spore Development. Dev. Cell 56, 36–51.e5. https://doi.org/10.1016/j.devcel.2020.12.006.

Molina-Quiroz, R.C., Lazinski, D.W., Camilli, A., and Levy, S.B.(2016). Transposon-Sequencing Analysis Unveils Novel Genes Involved in the Generation of Persister Cells in Uropathogenic Escherichia coli. Antimicrob. Agents Chemother. 60, 6907–6910. https://doi.org/10.1128/AAC.01617-16.

Moradali, M.F., Ghods, S., and Rehm, B.H.A.(2017). Pseudomonas aeruginosa Lifestyle: A Paradigm for Adaptation, Survival, and Persistence. Front. Cell. Infect. Microbiol. 7, 39. https://doi.org/10.3389/fcimb.2017.00039.

Osawa, K., Shigemura, K., Iguchi, A., Shirai, H., Imayama, T., Seto, K., Raharjo, D., Fujisawa, M., Osawa, R., and Shirakawa, T.(2013). Modulation of O-antigen chain length by the wzz gene in Escherichia coli O157 influences its sensitivities to serum complement. Microbiol Immunol 57, 616–623. https://doi.org/10.1111/1348-0421.12084.

Pacios, O., Blasco, L., Bleriot, I., Fernandez-Garcia, L., Ambroa, A., López, M., Bou, G., Cantón, R., Garcia-Contreras, R., Wood, T.K., et al. (2020). (p)ppGpp and Its Role in Bacterial Persistence: New Challenges. Antimicrob. Agents Chemother. 64, e01283–20. https://doi.org/10.1128/AAC.01283-20.

Pedersen, S.S., Kharazmi, A., Espersen, F., and Høiby, N. (1990). Pseudomonas aeruginosa alginate in cystic fibrosis sputum and the inflammatory response. Infect. Immun. 58, 3363–3368. https://doi.org/10.1128/iai.58.10.3363-3368.1990.

Phan, M.-D., Peters, K.M., Sarkar, S., Lukowski, S.W., Allsopp, L.P., Gomes Moriel, D., Achard, M.E.S., Totsika, M., Marshall, V.M., Upton, M., et al. (2013). The serum resistome of a globally disseminated multidrug resistant uropathogenic Escherichia coli clone. PLoS Genet. 9, e1003834. https://doi.org/10.1371/journal.pgen.1003834.

Pier, G.B., Coleman, F., Grout, M., Franklin, M., and Ohman, D.E.(2001). Role of alginate O acetylation in resistance of mucoid Pseudomonas aeruginosa to opsonic phagocytosis. Infect. Immun. 69, 1895–1901. https://doi.org/10.1128/IAI.69.3.1895-1901.2001.

Pont, S., Fraikin, N., Caspar, Y., Van Melderen, L., Attree, I., and Cretin, F.(2020). Bacterial behavior in human blood reveals complement evaders with some persister-like features. PLoS Pathog 16, e1008893. https://doi.org/10.1371/journal.ppat.1008893.

Pontes, M.H., and Groisman, E.A.(2019). Slow growth determines nonheritable antibiotic resistance in Salmonella enterica. Sci Signal 12. https://doi.org/10.1126/scisignal.aax3938.

Poulsen, B.E., Yang, R., Clatworthy, A.E., White, T., Osmulski, S.J., Li, L., Penaranda, C., Lander, E.S., Shoresh, N., and Hung, D.T.(2019). Defining the core essential genome of Pseudomonas aeruginosa. Proc. Natl. Acad. Sci. 116, 10072–10080. https://doi.org/10.1073/pnas.1900570116.

Priebe, G.P., Dean, C.R., Zaidi, T., Meluleni, G.J., Coleman, F.T., Coutinho, Y.S., Noto, M.J., Urban, T.A., Pier, G.B., and Goldberg, J.B.(2004). The galU Gene of Pseudomonas aeruginosa is required for corneal infection and efficient systemic spread following pneumonia but not for infection confined to the lung. Infect. Immun. 72, 4224–4232. https://doi.org/10.1128/IAI.72.7.4224-4232.2004.

Putrinš, M., Kogermann, K., Lukk, E., Lippus, M., Varik, V., and Tenson, T.(2015). Phenotypic Heterogeneity Enables Uropathogenic Escherichia coli To Evade Killing by Antibiotics and Serum Complement. Infect. Immun.

Qiu, D., Eisinger, V.M., Rowen, D.W., and Yu, H.D.(2007). Regulated proteolysis controls mucoid conversion in Pseudomonas aeruginosa. Proc. Natl. Acad. Sci. 104, 8107–8112. https://doi.org/10.1073/pnas.0702660104.

Samant, S., Lee, H., Ghassemi, M., Chen, J., Cook, J.L., Mankin, A.S., and Neyfakh, A.A.(2008). Nucleotide Biosynthesis Is Critical for Growth of Bacteria in Human Blood. PLoS Pathog. 4, e37. https://doi.org/10.1371/journal.ppat.0040037.

Sanchez-Larrayoz, A.F., Elhosseiny, N.M., Chevrette, M.G., Fu, Y., Giunta, P., Spallanzani, R.G., Ravi, K., Pier, G.B., Lory, S., and Maira-Litrán, T. (2017). Complexity of Complement Resistance Factors Expressed by Acinetobacter baumannii Needed for Survival in Human Serum. J. Immunol. Baltim. Md 1950 199, 2803–2814. https://doi.org/10.4049/jimmunol.1700877.

Shan, Y., Brown Gandt, A., Rowe, S.E., Deisinger, J.P., Conlon, B.P., and Lewis, K.(2017). ATP-Dependent Persister Formation in *Escherichia coli*. MBio 8. https://doi.org/10.1128/mBio.02267-16.

Short, F.L., Di Sario, G., Reichmann, N.T., Kleanthous, C., Parkhill, J., and Taylor, P.W.(2020). Genomic Profiling Reveals Distinct Routes To Complement Resistance in Klebsiella pneumoniae. Infect. Immun. 88. https://doi.org/10.1128/IAI.00043-20.

Skurnik, D., Roux, D., Aschard, H., Cattoir, V., Yoder-Himes, D., Lory, S., and Pier, G.B.(2013). A Comprehensive Analysis of In Vitro and In Vivo Genetic Fitness of Pseudomonas aeruginosa Using High-Throughput Sequencing of Transposon Libraries. PLoS Pathog. 9, e1003582. https://doi.org/10.1371/journal.ppat.1003582.

Trouillon, J., Sentausa, E., Ragno, M., Robert-Genthon, M., Lory, S., Attree, I., and Elsen, S.(2020). Species-specific recruitment of transcription factors dictates toxin expression. Nucleic Acids Res 48, 2388–2400. https://doi.org/10.1093/nar/gkz1232.

Ventre, I., Goodman, A.L., Vallet-Gely, I., Vasseur, P., Soscia, C., Molin, S., Bleves, S., Lazdunski, A., Lory, S., and Filloux, A.(2006). Multiple sensors control reciprocal expression of Pseudomonas aeruginosa regulatory RNA and virulence genes. Proc. Natl. Acad. Sci. 103, 171–176. https://doi.org/10.1073/pnas.0507407103.

Vitkauskienė, A., Skrodenienė, E., Dambrauskienė, A., Macas, A., and Sakalauskas, R.(2010). Pseudomonas aeruginosa bacteremia: resistance to antibiotics, risk factors, and patient mortality. Medicina (Mex.) 46, 490. https://doi.org/10.3390/medicina46070071.

Weber, B.S., De Jong, A.M., Guo, A.B.Y., Dharavath, S., French, S., Fiebig-Comyn, A.A., Coombes, B.K., Magolan, J., and Brown, E.D.(2020). Genetic and Chemical Screening in Human Blood Serum Reveals Unique Antibacterial Targets and Compounds against Klebsiella pneumoniae. Cell Rep. 32, 107927. https://doi.org/10.1016/j.celrep.2020.107927.

Winsor, G.L., Lam, D.K.W., Fleming, L., Lo, R., Whiteside, M.D., Yu, N.Y., Hancock, R.E.W., and Brinkman, F.S.L.(2011). Pseudomonas Genome Database: improved comparative analysis and population genomics capability for Pseudomonas genomes. Nucleic Acids Res. 39, D596–D600. https://doi.org/10.1093/nar/gkq869.

Winsor, G.L., Griffiths, E.J., Lo, R., Dhillon, B.K., Shay, J.A., and Brinkman, F.S.L.(2016). Enhanced annotations and features for comparing thousands of Pseudomonas genomes in the Pseudomonas genome database. Nucleic Acids Res. 44, D646–D653. https://doi.org/10.1093/nar/gkv1227.

Woong Park, S., Klotzsche, M., Wilson, D.J., Boshoff, H.I., Eoh, H., Manjunatha, U., Blumenthal, A., Rhee, K., Barry, C.E., Aldrich, C.C., et al. (2011). Evaluating the Sensitivity of Mycobacterium tuberculosis to Biotin Deprivation Using Regulated Gene Expression. PLoS Pathog. 7, e1002264. https://doi.org/10.1371/journal.ppat.1002264.

Wurtzel, O., Yoder-Himes, D.R., Han, K., Dandekar, A.A., Edelheit, S., Greenberg, E.P., Sorek, R., and Lory, S.(2012). The Single-Nucleotide Resolution Transcriptome of Pseudomonas aeruginosa Grown in Body Temperature. PLOS Pathog. 8, e1002945. https://doi.org/10.1371/journal.ppat.1002945.

Yin, Y., Damron, F.H., Withers, T.R., Pritchett, C.L., Wang, X., Schurr, M.J., and Yu, H.D.(2013). Expression of mucoid induction factor MucE is dependent upon the alternate sigma factor AlgU in Pseudomonas aeruginosa. BMC Microbiol. 13, 232. https://doi.org/10.1186/1471-2180-13-232.

Zhu, B., Green, S., Ge, X., Puccio, T., Nadhem, H., Ge, H., Bao, L., Kitten, T., and Xu, P.(2021). Genome-wide identification of Streptococcus sanguinis fitness genes in human serum and discovery of potential selective drug targets. Mol. Microbiol. 115. https://doi.org/10.1111/mmi.14629.

