## Supplementary material for "Molecular features underlying *Pseudomonas aeruginosa* persistence in human plasma": Suplemental information

### Supplementary information

#### Supplementary figures

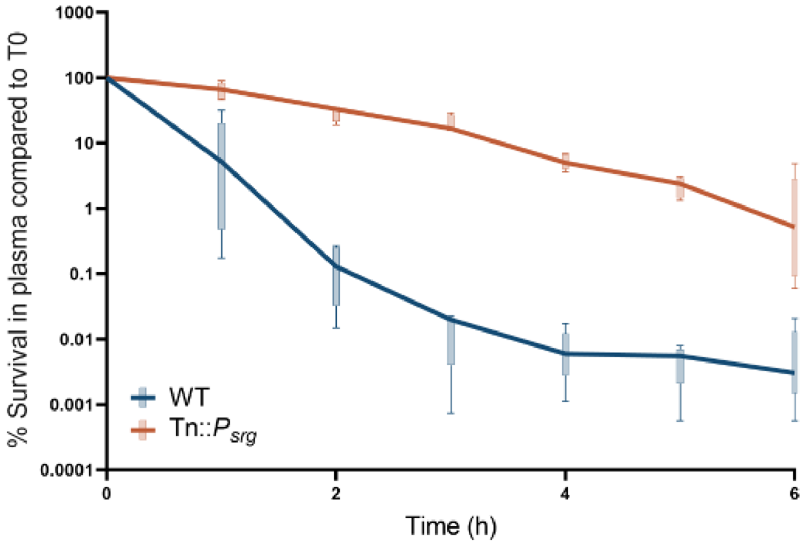

**Figure S1. Overexpression of *srg* operon leads to a population tolerant to plasma killing.** Survival kinetics of IHMA87 wild-type strain (dame data as presented in Fig. 2B) and of Tn::$P_{srg}$ in plasma over 6h incubation measured by CFU counting (n=5).

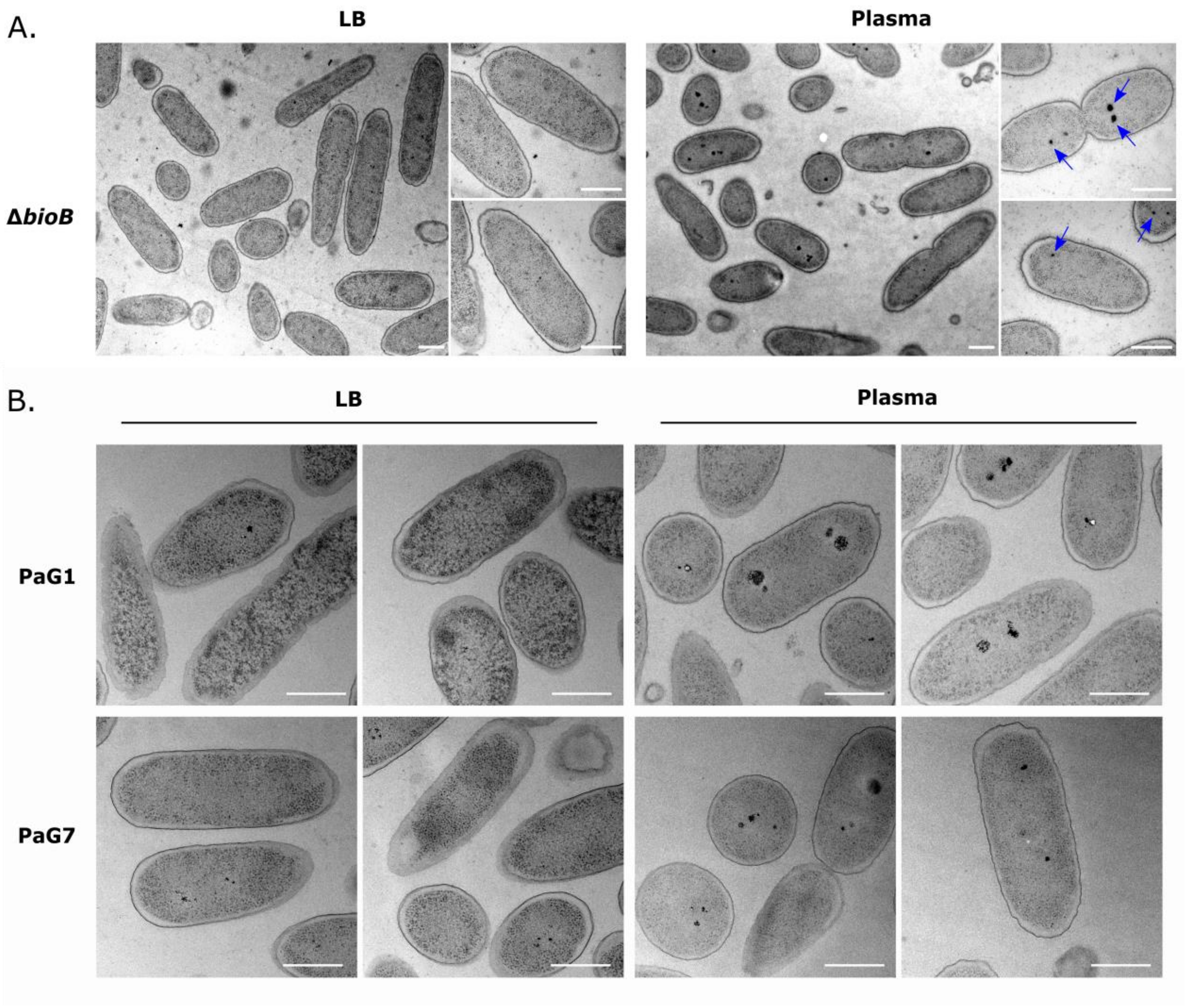

**Figure S2. Incubation in plasma triggers polyphosphate granules formation. A.** Transmission electron microscopy images of Δ*bioB* after growth in LB (left) or 1h-incubation in human plasma (right). Red and blue arrow show big- and small-sized granules respectively. **B.** Transmission electron microscopy images of two BSI isolates PaG1 and PaG7 after growth in LB (left) or 1h-incubation in human plasma (right). Scale bar = 500 nm.

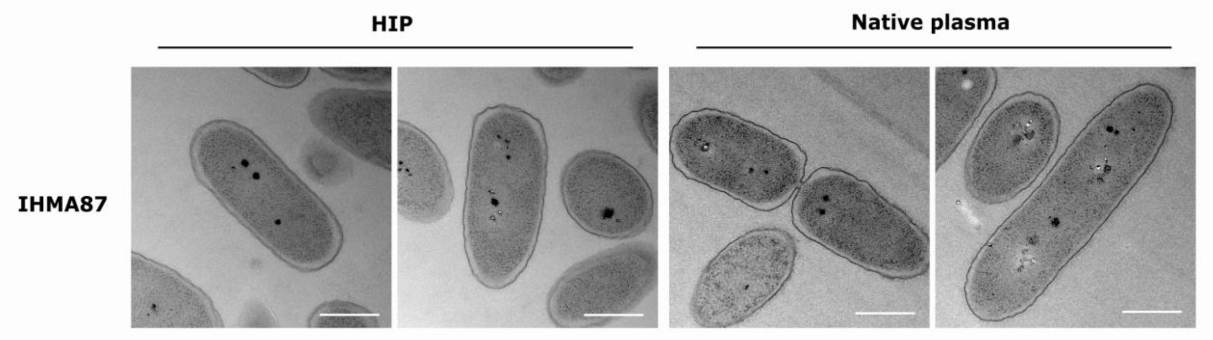

**Figure S3. Heat inactivated plasma triggers the formation of polyP granules.** Transmission electron microscopy images of IHMA87 after 1h-incubation in heat-inactivated plasma (HIP, left) or in native human plasma (right. Scale bar = 500 nm.

#### Supplementary tables

**Table 1S. Tn-seq data**

**Table 2S. RNA-seq differential gene expression data**

#### Table S3. Bacterial strains and plasmids

| Bacteria | Features | Reference/origin |
|---|---|---|
| ***Pseudomonas aeruginosa*** | | |
| IHMA879472/AZPAE15042 | Urinary isolate, plasma sensitive | IHMA[1] collection (1–3) |
| IHMA87Δ*bioB* | IHMA87 with *bioB* deletion | This work |
| IHMA87Δ*purD* | IHMA87 with *purD* deletion | This work |
| IHMA87Δ*algD* | IHMA87 with *algD* deletion | This work |
| IHMA87Δ*01134* | IHMA87 with *01134* deletion | This work |
| IHMA87 Tn::$P_{srg}$ | IHMA87 with mariner transposon inserted in the promoter of *srg* operon (Gm$^R$) | This work |
| IHMA87 Tn::$P_{srg}$ Δ*srgABC* | IHMA87 Tn::$P_{srg}$ with *srgABC* deletion (Gm$^R$) | This work |
| IHMA87 Tn::$P_{srg}$ Δ*srgA* | IHMA87 Tn::$P_{srg}$ with *srgA* deletion (Gm$^R$) | This work |
| IHMA87 Tn::$P_{srg}$ Δ*srgB* | IHMA87 Tn::$P_{srg}$ with *srgB* deletion (Gm$^R$) | This work |
| IHMA87 Tn::$P_{srg}$ Δ*srgC* | IHMA87 Tn::$P_{srg}$ with *srgC* deletion (Gm$^R$) | This work |
| IHMA87 Tn::$P_{srg}$ Δ*srgBC* | IHMA87 Tn::$P_{srg}$ with *srgBC* deletion (Gm$^R$) | This work |
| IHMA87 Tn::$P_{srg}$ Δ*algD* | IHMA87 Tn::$P_{srg}$ with *algD* deletion (Gm$^R$) | This work |
| IHMA87 Tn::$P_{srg}$ Δ*01134* | IHMA87 Tn::$P_{srg}$ with *01134* deletion (Gm$^R$) | This work |
| IHMA87 *ladS*::Tn | IHMA87 with mariner transposon inserted in *ladS* gene (Gm$^R$) | This work |
| IHMA87 *bioA*::Tn | IHMA87 with mariner transposon inserted in *bioA* gene (Gm$^R$) | This work |
| IHMA87 *bioB*::Tn | IHMA87 with mariner transposon inserted in *bioB* gene (Gm$^R$) | This work |
| IHMA87 *mucE*::Tn | IHMA87 with mariner transposon inserted in *mucE* gene (Gm$^R$) | This work |
| IHMA87 *mucA*::Tn | IHMA87 with mariner transposon inserted in *mucA* gene (Gm$^R$) | This work |
| IHMA87 *mucD*::Tn | IHMA87 with mariner transposon inserted in *mucD* gene (Gm$^R$) | This work |
| IHMA87 *pprB*::Tn | IHMA87 with mariner transposon inserted in *pprB* gene (Gm$^R$) | This work |
| IHMA87 *mucB*::Tn | IHMA87 with mariner transposon inserted in *mucB* gene (Gm$^R$) | This work |
| PaG1 | Bloodstream isolate, O8, infected organ | (2) |
| PaG7 | Bloodstream isolate, O1, contaminated catheter | (2) |
| ***Escherichia coli*** | | |
| DH5α | Laboratory strain | Lab collection |
| TOP10 | Cloning strain | Invitrogen |
| **Plasmids** | | |
| pBTK24 | Plasmid with Himar-1 mariner transposon and C9 transposase (Amp$^R$, Gm$^R$) | (4) |
| pRK600 | Helper plasmid with conjugative properties (Cm$^R$) | (5) |
| pEXG2 | Allelic exchange vector (Gm$^R$), *sacB* | (6) |
| pEX18Tc | Allelic exchange vector (Tc$^R$), *sacB* | (7) |
| pEX100T | Allelic exchange vector (Cb$^R$), *sacB* | (8) |
| pminiCTX-*pX2*-GFP | pminiCTX harboring the *gfp* gene under the control of a constitutive promoter (*attP* site, Tc$^R$) | (9) |

| | | |
|---|---|---|
| pminiCTX-*Psrg-lacZ* | pminiCTX-*lacZ* harboring the srg promoter fused to *lacZ* reporter gene (*attP* site, Tc$^R$) | This work |
| pEXG2-mut-*bioB* | pEXG2 carrying DNA fragment for *bioB* deletion obtained by SLIC (Gm$^R$) | This work |
| pEXG2-mut-*purD* | pEXG2 carrying DNA fragment for *purD* deletion obtained by SLIC (Gm$^R$) | This work |
| pUC57-mut-*srgABC* | pUC57 carrying the synthetic DNA fragment for *srgABC* deletion in Tn::$P_{srg}$ (Genewiz) | This work |
| pEX18Tc-mut-*srgABC* | pEX18Tc carrying DNA fragment for srgABC deletion in Tn::$P_{srg}$ mutant obtained by SLIC (Tc$^R$) | This work |
| *pEX18Tc-mut-01134* | pEX18Tc carrying DNA fragment for IHMA87_01134 deletion obtained by SLIC (Tc$^R$) | This work |
| pEX100T-mut-*srgA* | pEX100T carrying DNA fragment for *srgA* deletion obtained by SLIC (Cb$^R$) | This work |
| pEX100T-mut-*srgB* | pEX100T carrying DNA fragment for *srgB* deletion obtained by SLIC (Cb$^R$) | This work |
| pEX100T-mut-*srgC* | pEX100T carrying DNA fragment for *srgC* deletion obtained by SLIC (Cb$^R$) | This work |
| pEX100T-mut-*srgAB* | pEX100T carrying DNA fragment for *srgAB* deletion obtained by SLIC (Cb$^R$) | This work |
| pEX100T-mut-*srgBC* | pEX100T carrying DNA fragment for *srgBC* deletion obtained by SLIC (Cb$^R$) | This work |
| pEX100T-mut-*algD* | pEX100T carrying DNA fragment for *algD* deletion obtained by SLIC (Cb$^R$) | This work |

[1] International Health Management Association, USA

#### Table S4. Oligonucleotides used for PCRs

| **Primers** | **Sequence (5'-3')** | |
|---|---|---|
| pEXG2-mut-*bioB*-sF1 | **GGTCGACTCTAGAGGATCCCC**GTTCGG ACCCCTCGAACATCG | *bioB* deletion |
| pEXG2-mut-*bioB*-sR1 | GGTGGCGACGGCTGCGGAT | *bioB* deletion |
| pEXG2-mut-*bioB*-sF2 | **GCGCATCCGCAGCCGTCGCCACC**CAGC TGTTCTATAACGCCGCCT | *bioB* deletion |
| pEXG2-mut-*bioB*-sR2 | **ACCGAATTCGAGCTCGAGCCC**CTTGCC GACCAGCGCGGTGA | *bioB* deletion |
| F0-mut-*bioB* | GCGAGCAGCCGTTCGGCG | PCR verification for *bioB* deletion |
| R0-mut-*bioB* | TTGAGGCGGTCCTCCAGCA | PCR verification for *bioB* deletion |
| pEXG2-mut-*purD*-sF1 | **GGTCGACTCTAGAGGATCCCC**CAGGAG ATCCACGACCTGATCT | *purD* deletion |
| pEXG2-mut-*purD*-sR1 | ACCGCCGCTGCCGATGATGA | *purD* deletion |
| pEXG2-mut-*purD*-sF2 | **TACTCATCATCGGCAGCGGCGGT**GAGC GCGGCGAGTCCTGAC | *purD* deletion |
| pEXG2-mut-*purD*-sR2 | **ACCGAATTCGAGCTCGAGCCC**TCGCGG ACCAGCTGGCCGT | *purD* deletion |
| F0-mut-*purD* | CTGAAGATCGTCACTCGCCG | PCR verification for *purD* deletion |
| R0-mut-*purD* | GACTCGCCGGTGTGCTGGT | PCR verification for *purD* deletion |
| pEX18Tc-mut-*srgABC*-sF1 | **GTCGACTCTAGAGGATCCC**CGGATCCC CGACATTCGGCTA | Transfer DNA fragment from pUC57 to pEX18Tc for *srgABC* deletion in Tn::$P_{srg}$ |
| pEX18Tc-mut-*srgABC*-sR1 | **CGAATTCGAGCTCGGTACCC**AAGCTTT GATCTACGTGCAAGC | |

| | | |
|---|---|---|
| F0–mut-*srgABC* | GCCTCGCCGACCTCTACA | PCR verification for *srgA, srgB, srgC* and *srgABC* deletion |
| RO-Tn::$P_{srg}$-mut-*srgABC* | GCTTGCTGCCTTCGACCAAG | PCR verification for *srgA, srgB, srgC* and *srgABC* deletion |
| pEX18Tc-mut-01134-sF1 | **GTCGACTCTAGAGGATCCCC**CACCTCG GTGTCCACGCTGC | IHMA87_01134 deletion |
| pEX18Tc-mut-01134-sR1 | CGACGAGCATCGACTTGTTCAC | IHMA87_01134 deletion |
| pEX18Tc-mut-01134-sF2 | **TGAACAAGTCGATGCTCGTC**GACAAGA ACGGTCGGCTGGT | IHMA87_01134 deletion |
| pEX18Tc-mut-01134-sR2 | **CGAATTCGAGCTCGGTACCC**ATGTTCC GCAGCCTGGTCGG | IHMA87_01134 deletion |
| F0-mut-01134 | CTGCTGTACGTCGGCTTCCG | PCR verification for IHMA87_01134 deletion |
| R0-mut-01134 | CTGGATGGCACCACCAACTTC | PCR verification for IHMA87_01134 deletion |
| pEX100T-mut-*srgA*-sF1 | **ACCCTGTTATCCCTACCC**GTGATCTACG TGCAAGCAGA | *srgA* and *srgAB* deletion in Tn::$P_{srg}$ |
| pEX100T-mut-*srgA*-sR1 | CAGGTTGCGTGAGCTGCTCA | *srgA* and *srgAB* deletion in Tn::$P_{srg}$ |
| pEX100T-mut-*srgA*-sF2 | **TGAGCAGCTCACGCAACCTGTGA**CCTG GACATTCTGACGAGGTA | *srgA* deletion in Tn::$P_{srg}$ |
| pEX100T-mut-*srgA*-sR2 | **GGATAACAGGGTAATCCC**CACCAGCGC AAGCAGCAGC | *srgA* deletion in Tn::$P_{srg}$ |
| pEX100T-mut-*srgB*-sF1 | **ACCCTGTTATCCCTACCC**ATGGAGGTGA ACATGAGCAGC | *srgB* and *srgBC* deletion in Tn::$P_{srg}$ |
| pEX100T-mut-*srgB*-sR1 | AGCCAGAATCCACATGTTTC | *srgB* and *srgBC* deletion in Tn::$P_{srg}$ |
| pEX100T-mut-*srgB*-sF2 | **GAAACATGTGGATTCTGGCTTGA**ACGG AAGCTCGGCGCCACT | *srgB* deletion in Tn::$P_{srg}$ |
| pEX100T-mut-*srgB*-sR2 | **GGATAACAGGGTAATCCC**CGTGTTGAC TCACGTCGGAC | *srgB* deletion in Tn::$P_{srg}$ |
| pEX100T-mut-*srgC*-sF1 | **ACCCTGTTATCCCTACCC**CGACGAACA GGCCTGGACAT | *srgC* deletion in Tn::$P_{srg}$ |
| pEX100T-mut-*srgC*-sR1 | AAGAATGACGGCGGATCGTGA | *srgC* deletion in Tn::$P_{srg}$ |
| pEX100T-mut-*srgC*-sF2 | **TCACGATCCGCCGTCATTCTTTGA**GATC CCAGCCTGGACTGATC | *srgC* deletion in Tn::$P_{srg}$ |
| pEX100T-mut-*srgC*-sR2 | **GGATAACAGGGTAATCCCC**CACTCGAA GCCGACATTCG | *srgC* deletion in Tn::$P_{srg}$ |
| pEX100T-mut-*srgAB*-sF2 | **TGAGCAGCTCACGCAACCTGTGA**ACGG AAGCTCGGCGCCACT | *srgAB* deletion in Tn::$P_{srg}$ |
| pEX100T-mut-*srgAB*-sR2 | **GGATAACAGGGTAATCCC**CGTGTTGAC TCACGTCGGAC | *srgAB* deletion in Tn::$P_{srg}$ |
| pEX100T-mut-*srgBC*-sF2 | **GAAACATGTGGATTCTGGCTTGA**GATC CCAGCCTGGACTGATC | *srgBC* deletion in Tn::$P_{srg}$ |
| pEX100T-mut-*srgBC*-sR2 | **GGATAACAGGGTAATCCCC**CACTCGAA GCCGACATTCG | *srgBC* deletion in Tn::$P_{srg}$ |
| pEX100T-mut-*algD*-sF1 | **ACCCTGTTATCCCTACCC**CGGAAACGC CATCAAGTTGGTA | *algD* deletion |
| pEX100T-mut-*algD*-sR1 | GCCAGCACATACCGCACCGA | *algD* deletion |
| pEX100T-mut-*algD*-sF2 | **TCGGTGCGGTATGTGCTGGC**CAGGCCG AGGGCATCTGCT | *algD* deletion |
| pEX100T-mut-*algD*-sR2 | **GGATAACAGGGTAATCCC**GCTGTCGAT GGCTTCGCGGA | *algD* deletion |
| F0-mut-*algD* | CTCGTGGCGAATAGGCCTAC | PCR verification for *algD* deletion |
| R0-mut-*algD* | GCGAAGCGCAGCTTGTGCC | PCR verification for *algD* deletion |

| Primers for Tn-seq | |
|---|---|
| Short adpator | TACCACGACCA-NH2 |
| Long adaptor | GTGACTGGAGTTCAGACGTGTGCTCTTC CGATCTGGTCGTGGTAT |
| PCR1 Tn-specific | CACAGGAAACAGGACTCTAGAGG |
| PCR2 adaptor complementary | GTGACTGGAGTTCAGACGTGTG |
| P5+ Illumina | AATGATACGGCGACCACCGAGATCTACA CTCTTTCCCTACACGACGCTCTTCCGAT CTCTAGAGACCGGGGACTTATCAGC |
| P7-index | CAAGCAGAAGACGGCATACGAGAT**CGT GAT** |

#### Supplementary references

1. Kos VN, Deraspe M, McLaughlin RE, Whiteaker JD, Roy PH, Alm RA, et al. The resistome of Pseudomonas aeruginosa in relationship to phenotypic susceptibility. Antimicrob Agents Chemother. 2015 Jan;59(1):427–36.

2. Pont S, Fraikin N, Caspar Y, Van Melderen L, Attree I, Cretin F. Bacterial behavior in human blood reveals complement evaders with some persister-like features. PLoS Pathog. 2020 Dec;16(12):e1008893.

3. Trouillon J, Sentausa E, Ragno M, Robert-Genthon M, Lory S, Attree I, et al. Species-specific recruitment of transcription factors dictates toxin expression. Nucleic Acids Res. 2020 Mar 18;48(5):2388–400.

4. Kulasekara HD, Ventre I, Kulasekara BR, Lazdunski A, Filloux A, Lory S. A novel two-component system controls the expression of Pseudomonas aeruginosa fimbrial cup genes. Mol Microbiol. 2005;55(2):368–80.

5. Kessler B, de Lorenzo V, Timmis KN. A general system to integratelacZ fusions into the chromosomes of gram-negative eubacteria: regulation of thePm promoter of theTOL plasmid studied with all controlling elements in monocopy. Mol Gen Genet MGG. 1992 May;233(1–2):293–301.

6. Rietsch A, Vallet-Gely I, Dove SL, Mekalanos JJ. ExsE, a secreted regulator of type III secretion genes in Pseudomonas aeruginosa. Proc Natl Acad Sci. 2005 May 31;102(22):8006–11.

7. Li S, Li X, Zhao H, Cai B. Physiological role of the novel salicylaldehyde dehydrogenase NahV in mineralization of naphthalene by Pseudomonas putida ND6. Microbiol Res. 2011 Dec 20;166(8):643–53.

8. Schweizer HP, Hoang TT. An improved system for gene replacement and xylE fusion analysis in Pseudomonas aeruginosa. Gene. 1995 Jan 1;158(1):15–22.

9. Thibault J, Faudry E, Ebel C, Attree I, Elsen S. Anti-activator ExsD forms a 1:1 complex with ExsA to inhibit transcription of type III secretion operons. J Biol Chem. 2009 Jun 5;284(23):15762–70.
